# Impaired loading following denervation elicits dynamic, differential growth trajectories of musculoskeletal tissues

**DOI:** 10.64898/2026.10.26.761680

**Authors:** Talayah Johnson, Natalie L. Fogarty, Grey Hallstrom, Destiny Miller, Madison Dennis, Alisia Lin, Xi Jiang, Jiaqi Xiang, Joel D. Boerckel, Eiki Koyama, Lin Han, Josh R. Baxter, Robert L. Mauck, Nathaniel A. Dyment

## Abstract

Physical forces dictate the function and maintenance of adult musculoskeletal tissues. Yet, the impact of biophysical cues on the growth and maturation of these tissues, especially dense connective tissues, remains unclear. Here, we investigated the effects of neonatal mechanical loading on early postnatal growth of the musculoskeletal tissues of the lower limb, with particular focus on the Achilles tendon. We performed sciatic nerve resection (SNR) on one-day-old mice to denervate the hamstrings and distal hindlimb muscles. Sciatic Nerve Resection resulted in severe and sustained alterations in gait, accompanied by an altered transcriptional landscape in distal musculoskeletal tissues, with marked reductions in bone and tendon growth and smaller, fibrotic hindlimb musculature. Sciatic nerve resection impaired cumulative growth in the Achilles tendon, shifting cell proliferation and matrix-driven growth trajectories and altering mechanotransduction gene programs toward inflammation and matrix catabolism with age. This work highlights the essential role of mechanical loading in driving mechanosignaling and postnatal growth trajectories in musculoskeletal tissues of the developing limb, with implications for overall musculoskeletal health in adulthood.

## INTRODUCTION

Mechanical forces are vital for musculoskeletal tissue formation and maintenance, influencing how tissues grow, adapt, and maintain function throughout life (*1–6*). Declines in physical activity, whether due to neurological disorders or to increased sedentary behavior, are linked to impairments in musculoskeletal health, including poorer bone quality and an increased risk of degenerative diseases such as osteoarthritis, underscoring the importance of mechanical loading throughout the lifespan (*7–9*). Even though it is well established that mechanical forces are critical to the overall health and function of musculoskeletal tissues, we still have a limited understanding of the roles of mechanical cues during the postnatal growth of dense connective tissues, a critical window for establishing the structural specialization required to meet tissue-specific functional demands. This knowledge gap limits our ability to sufficiently treat patients in need, particularly those suffering from neurological disorders (e.g., cerebral palsy). Additionally, in a society where children are becoming less physically active and more sedentary, it is important to consider the long-term consequences of reduced mechanical loading during this developmental period, particularly for dense connective tissues, which have minimal turnover in adulthood (*10*).

One such tissue that is highly sensitive to mechanical cues is the tendon. It is postulated that mechanical forces generated during embryonic and postnatal stages provide the cues that guide proper tendon formation and maturation. For example, embryonic immobilization in utero impairs tendon growth and maturation, impairing tissue function (*11–13*). The structural and functional properties of tendons, characterized by load-bearing extracellular matrix (ECM) dominated by highly aligned collagen I fibers, are established through tightly regulated processes during embryonic and postnatal growth (*14–17*). *In utero*, tendon progenitors self-organize into linear arrays, even before substantial ECM deposition (*18*). These cells transduce mechanical cues through the actin cytoskeleton in response to applied forces. Embryonic studies provide evidence that mechanical loading, particularly from muscle contraction, is required for tendon growth (*11*, *19–22*). For example, embryonic mice lacking skeletal muscle or muscle contractility (i.e., *mdg*/*mdg* mutants) exhibit impaired development, with decreased tendon cross-sectional area, reinforcing the idea that intrinsic mechanical forces drive tendon morphogenesis (*23*, *24*). Similarly, these models also develop severe skeletal abnormalities, highlighting the necessity of fetal movements for the development of the limb mesenchyme (*25–27*). While embryonic studies provide strong evidence that loading regulates the formation and development of dense connective tissues of the limb, *in vivo* studies investigating its role in early postnatal growth remain limited.

A critical shift in tendon growth occurs in the neonatal period. This includes a marked increase in tendon length and cross-sectional area over the first 1–2 weeks after birth, corresponding with high levels of cell proliferation (*25*, *26*). After postnatal day 14 (P14), proliferation dramatically declines, while substantial growth continues, mediated in this period by ECM synthesis and assembly. This transition from proliferation-driven to matrix-driven growth defines a critical developmental window. During this time, tendon cells align into linear arrays with established cell-cell junctions and then synthesize substantial ECM between these arrays, which is essential for mechanical strength and longitudinal growth (*15*, *25*)(*15*, *25*, *27*). This developmental window is particularly significant, as it coincides with the onset of weight-bearing ambulation, a stage at which mechanical forces acting on these tissues increase markedly. Therefore, mechanical loading may play a critical role in the shift from proliferation-driven growth to ECM-driven growth in neonatal tendons; however, the underlying mechanisms and regulatory factors governing this transformation are not yet defined.

In animal models, several methods have been employed to perturb mechanical loading, including botulinum toxin injections, tail suspension, tendon injury, and immobilization via casting (*28*, *29*). For example, enthesis maturation and increases in supraspinatus tendon properties were hindered following postnatal Botox injection (*30–32*). However, many of these models present technical challenges when applied at very early postnatal stages. A promising alternative for this age range is sciatic nerve resection (SNR) (*33–35*), which results in prolonged, persistent gait deficits and alterations in hindlimb loading bearing utilization, making it a valuable model for studying the effects of mechanical unloading across the lifespan (*33*, *34*). Previous work from our group and others has shown that SNR disrupts gait, reduces tibial growth, and alters growth plate structure (*33*, *36*). In addition, SNR reduces mechanical loading without directly injuring mesenchymal tissues, allowing the effects of chronic mechanical loading impairment to be studied in a physiologically relevant context.

Understanding how tendons, muscles, and bones respond to altered loading during this critical window may reveal key mechanotransduction pathways governing growth and long-term function. To investigate this, we employed surgical sciatic nerve resection (SNR) at postnatal day 1 (P1) to impair mechanical loading of the hindlimb during early postnatal development. Our findings reveal that sustained gait disruptions following SNR elicit dynamic, specific shifts in the growth trajectories of several tissues in the distal hindlimb. We observed a shift in muscle toward fibrotic remodeling, impaired bone growth with notable alterations in periosteal cell populations, and compromised tendon growth and maturation. Notably, although tendon growth was impaired, the growth period was prolonged, with later stages suggesting a progression to a pro-inflammatory and catabolic state, likely due to impaired matrix formation that was insufficient to withstand the demands of activities of daily living in adulthood.

## RESULTS

### Sciatic nerve resection (SNR) in the early postnatal period results in sustained gait deficits

Following the sciatic nerve resection (SNR) procedure on postnatal day 1 (P1), mice recovered well and were reintroduced to their litters, as in our previous study (*33*). In the first few days after surgery, the animals dragged the operated limb (**Supplemental video 1**). As they began to ambulate (∼P7), they adapted their gait to compensate for the impaired sciatic nerve function (**Supplemental videos 2-4**). To monitor these changes, we used video gait analysis in a longitudinal cohort of mice at multiple postnatal time points to investigate changes in joint kinematics as a consequence of SNR, with a particular focus on the distal limb to complement our previously reported gait parameters (*33*). SNR limbs exhibited persistent gait abnormalities compared to contralateral limbs. Toe spread was significantly reduced at all time points (**Fig. 1B**, p < 0.01), a hallmark response to sciatic denervation (*37*, *38*). In addition, the SNR limb showed marked increases at all time points in ankle dorsiflexion at paw strike and at lift-off, indicating collapse of the ankle at strike and diminished push-off strength, respectively (**Fig. 1C**, p < 0.01). Similarly, hock height at paw strike and lift off was significantly lower in the SNR limb throughout the duration of the study (**Fig. 1C**, p < 0.01). These deficits persisted through all developmental stages assessed in this study, indicating a lack of nerve regeneration and sustained alterations in gait.

**Figure 1:**
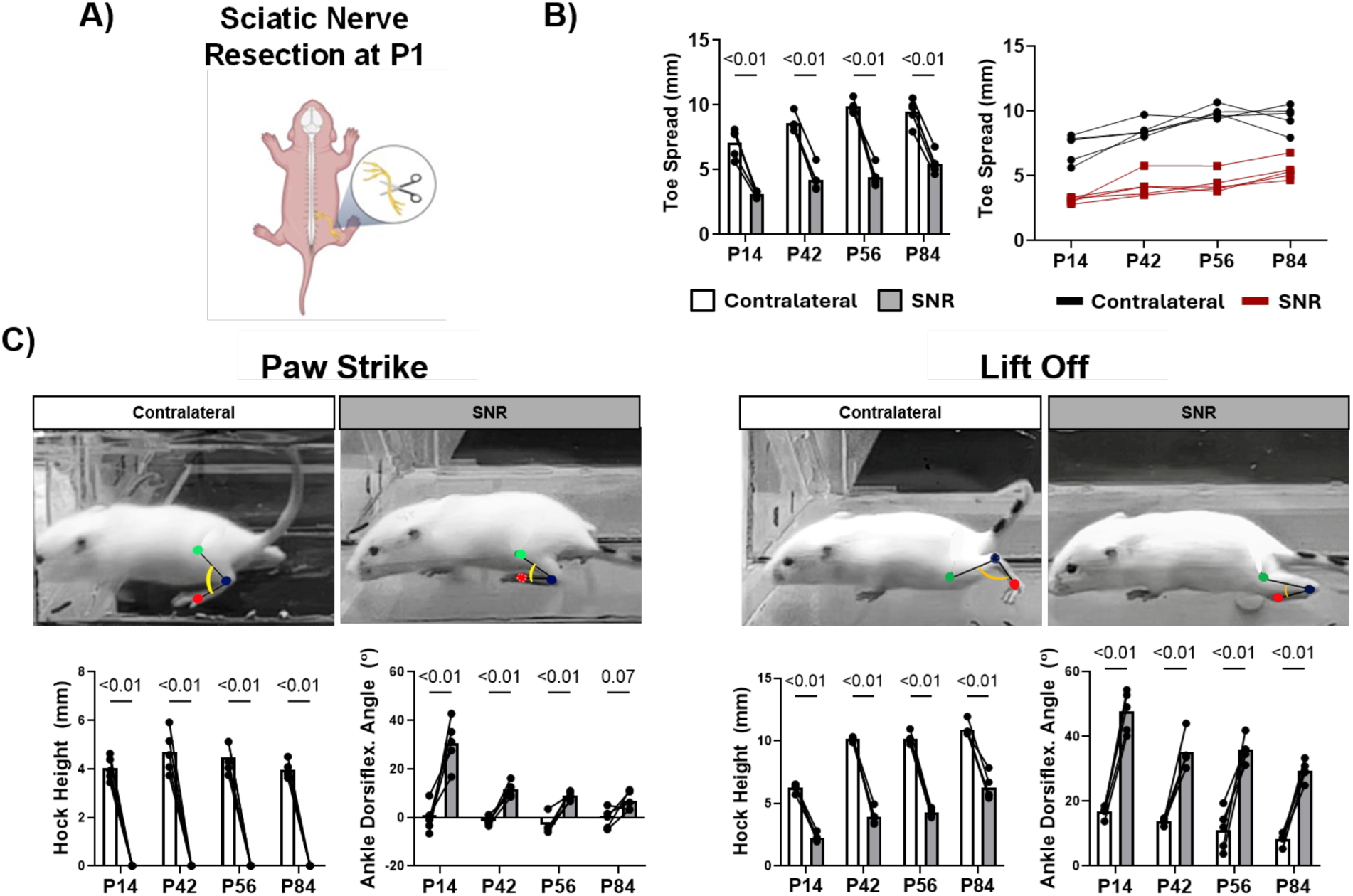
Longitudinal gait alterations following SNR performed on P1 mice. Video gait analysis following A) unilateral sciatic nerve resection (SNR) procedure performed at postnatal day 1 (P1). B) SNR limbs demonstrated a significant decrease in toe spread compared to the contralateral control limbs at all time points. C) Gait analysis images with labeled knee (green), ankle (blue), and metatarsophalangeal (red) joints. Changes in hock height and ankle dorsiflexion were apparent during both paw strike and lift off. SNR yielded decreased hock height and increased ankle dorsiflexion at all timepoints (p < 0.01, n=5).

### SNR drives broad transcriptional changes in muscle, nerve, bone, and tendon tissues

To provide spatial context for cellular changes across tissues affected by the altered loading from SNR, we performed Xenium spatial transcriptomic profiling to establish gene expression patterns at single-cell resolution. This was done on both the SNR and contralateral limbs at P14 (**Fig. 2**), a time point at which uninjured animals ambulate consistently and there is a significant impairment in gait in the SNR limb. Sagittal sections encompassing the entire Achilles tendon and portions of the adjacent muscle, nerves, fat pad, and bone were analyzed (**Fig. 2A, D and S1**). After quality filtering, a total of 299,663 cells were analyzed, including 141,006 contralateral cells and 158,657 SNR cells. Unbiased Leiden clustering identified 9 transcriptionally distinct clusters, which were annotated based on differentially expressed genes (DEGs), known cell-type markers, and spatial localization (**Fig. 2B**). The top-ranked DEGs for each cluster, determined by log_2_-fold change and enrichment, are summarized in **Figure S1A**. A UMAP projection, colored by treatment, revealed that while all clusters contained cells from both SNR and contralateral limbs, several populations showed marked compositional shifts following denervation, most notably nerve cells, skeletal muscle cells, osteochondral cells, and tendon internal fibroblasts (IFs) (dashed outlines in **Fig. 2C** and heatmap in **Fig. 2E**).

**Figure 2:**
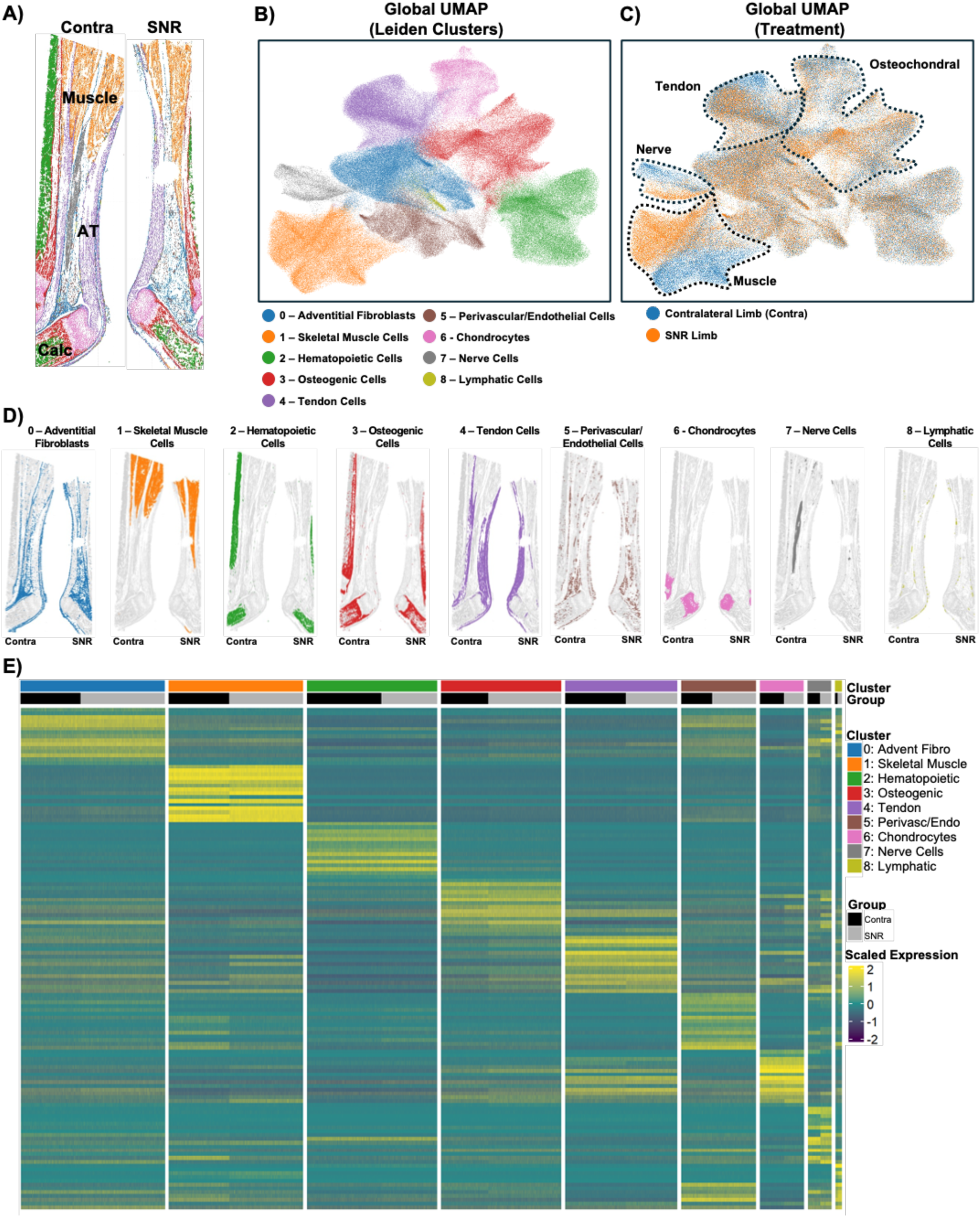
SNR alters the transcriptional landscape and cellular composition of nerve, muscle, bone, and tendon. Xenium spatial transcriptomics was performed on P14 hindlimb sections using a 479-gene probe panel (n = 3 mice/treatment, 4 sections/mouse). A) Spatial mapping of all Leiden clusters on the sections, illustrating their anatomical location in each tissue. B) Global UMAP showing 9 transcriptionally distinct populations following Leiden clustering. C) UMAP colored by surgical treatment with Leiden clusters affected by surgical treatment (muscle, nerve, tendon, and osteochondral) outlined. (AT: Achilles tendon; Calc: Calcaneus). D) Spatial mapping of each individual Leiden cluster. E) Heatmap of scaled gene expression across all Leiden clusters and treatments.

Cluster 7 in the global analysis (**Fig. 2B**) was identified as a nerve cell population based on location and expression of markers including *Kcna1, Plp1, Prx,* and *Ndgr1*. UMAP projection colored by treatment revealed a marked shift in this population following surgical treatment (**Fig. 2C**). SNR decreased the expression of mature Schwann cell-associated genes such as *Pmp22*, *Prx*, and *Ndrg1*, while increasing the expression of genes linked to axonal repair and remodeling (*Tubb3, Gap43, Runx2*) (*39–41*) in these cells (**Fig. S1B**).

### Skeletal muscle becomes fibrotic following SNR

Cluster 1 from the global analysis was classified as skeletal muscle based on canonical muscle gene expression, including *Myf6*, *Myoz1, Actn3*, *Trdn, Pvalb*, *Ttn, Des, Eno3, and Hrc*. Although represented as a single population in the global analysis, SNR and contralateral cells occupied largely distinct regions of the skeletal muscle cluster, indicating a pronounced treatment-associated shift in transcriptional state (**Fig. 2C**). Therefore, we reanalyzed these muscle associated cells, which resulted in 6 subclusters (**Fig. 3A and S2**). Three subclusters mapped spatially to myogenic cells, with two clusters that were predominant in the contralateral limb (Cluster 2: mostly in gastrocnemius and Cluster 4: mostly in soleus), whereas Cluster 0 was almost exclusively in the SNR muscle (**Fig. 3A-C**). The cluster-marker heatmap further revealed distinct transcriptional identities across these and other subclusters between contralateral and SNR cells (**Fig. 3E**). In addition, the density of fibroblasts and macrophages were increased in the SNR muscle (**Fig. 3D**). When comparing myogenic markers differentially expressed in the contralateral and SNR myogenic cells (Clusters 0, 2, 4), the SNR cells were downregulated for *Myoz2*, *Ppargc1a*, and *Eno3* and upregulated for *Myf6*, *Tnnc1*, and *Mustn1* (**Fig. 3F**). The SNR muscle consisted of increased centrally located nuclei and phalloidin-negative myofibers with elevated staining for the fibroblast marker PDGFRα (**Fig. 3G**), indicating an actively remodeling muscle. Together, these findings indicate that SNR induced fibrotic remodeling in the skeletal muscle.

**Figure 3:**
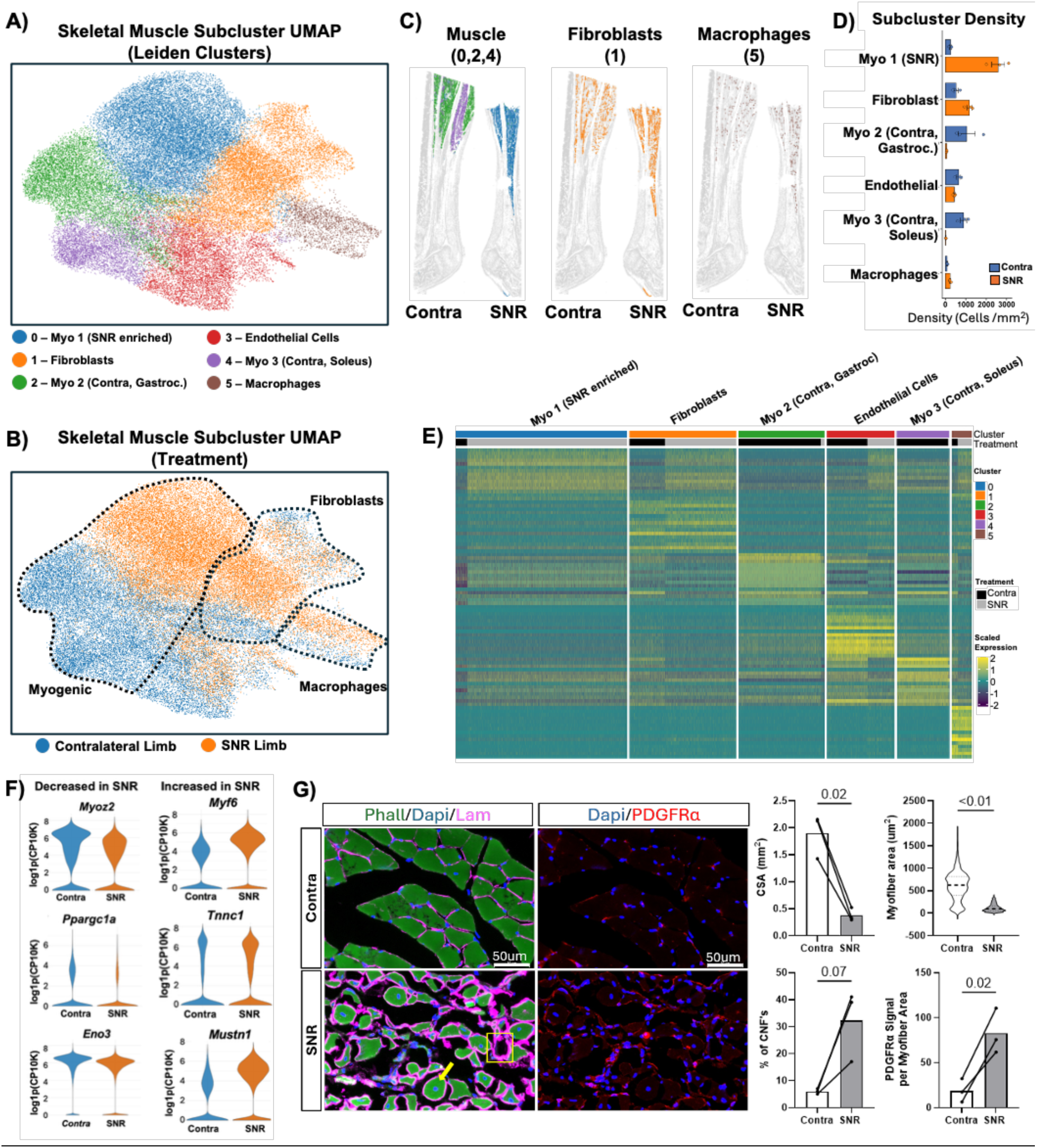
SNR causes muscle fibrosis and a phenotypic shift in myofibers. A) UMAP of reanalysis of Leiden cluster 1 (skeletal muscle), revealing six subclusters: three myogenic populations [Myo 1 (SNR-enriched), Myo 2 (Contralateral, gastrocnemius-enriched), and Myo 3 (contralateral, soleus-enriched)], fibroblasts, endothelial cells, and macrophages. B) Skeletal muscle UMAP colored by treatment, revealing treatment-associated changes in transcriptional profiles (dashed outlines highlight shifts in the myogenic, fibroblast, and macrophage populations). C) Spatial mapping of muscle cells, fibroblasts, and macrophages in contralateral and SNR sections. D) Cell density of each subcluster within the muscle. E) Heatmap of scaled gene expression across skeletal muscle subclusters and treatments. F) Differential expression of genes between SNR and contralateral myogenic cells (pooled Myogenic 1-3 clusters). G) Immunofluorescence images (n = 3) of Phalloidin, anti-laminin, anti-PDGFRα, and DAPI staining in contralateral and SNR muscle at P42. Quantification of gastrocnemius cross-sectional area, individual myofiber area, percentage of centrally nucleated fibers (CNFs), and PDGFRα fluorescence normalized to total myofiber area. Arrow indicates a centrally nucleated fiber, and box highlights a phalloidin-negative fiber. Scale bars = 50 µm.

### SNR compromises bone growth and development in the distal limb

We previously reported a marked reduction in the size of the hindlimb musculature following SNR (*33*), in combination with the gait alterations. We hypothesized that this would impair bone growth due to altered loading. Therefore, we investigated overall tibial growth, including trabecular and cortical bone parameters. Consistent with previous studies (*33*, *35*, *36*), neonatal SNR significantly reduced bone growth, as evidenced by reduced tibia length at P42 (**Fig. 4A**, p < 0.01). Significant differences in trabecular bone were apparent at P42 and P84 as well: specifically, trabecular bone volume fraction (BV/TV) and bone mineral density (BMD) (p<0.01, **Fig. 4B**) were reduced in the SNR limbs, among other parameters (**Fig. S3**). Similarly, cortical total area and cortical thickness (p<0.05, **Fig. 4C)** were reduced in the SNR limb compared to the contralateral limb at P14, P42, and P84 (additional parameters in **Fig. S3**).

**Figure 4:**
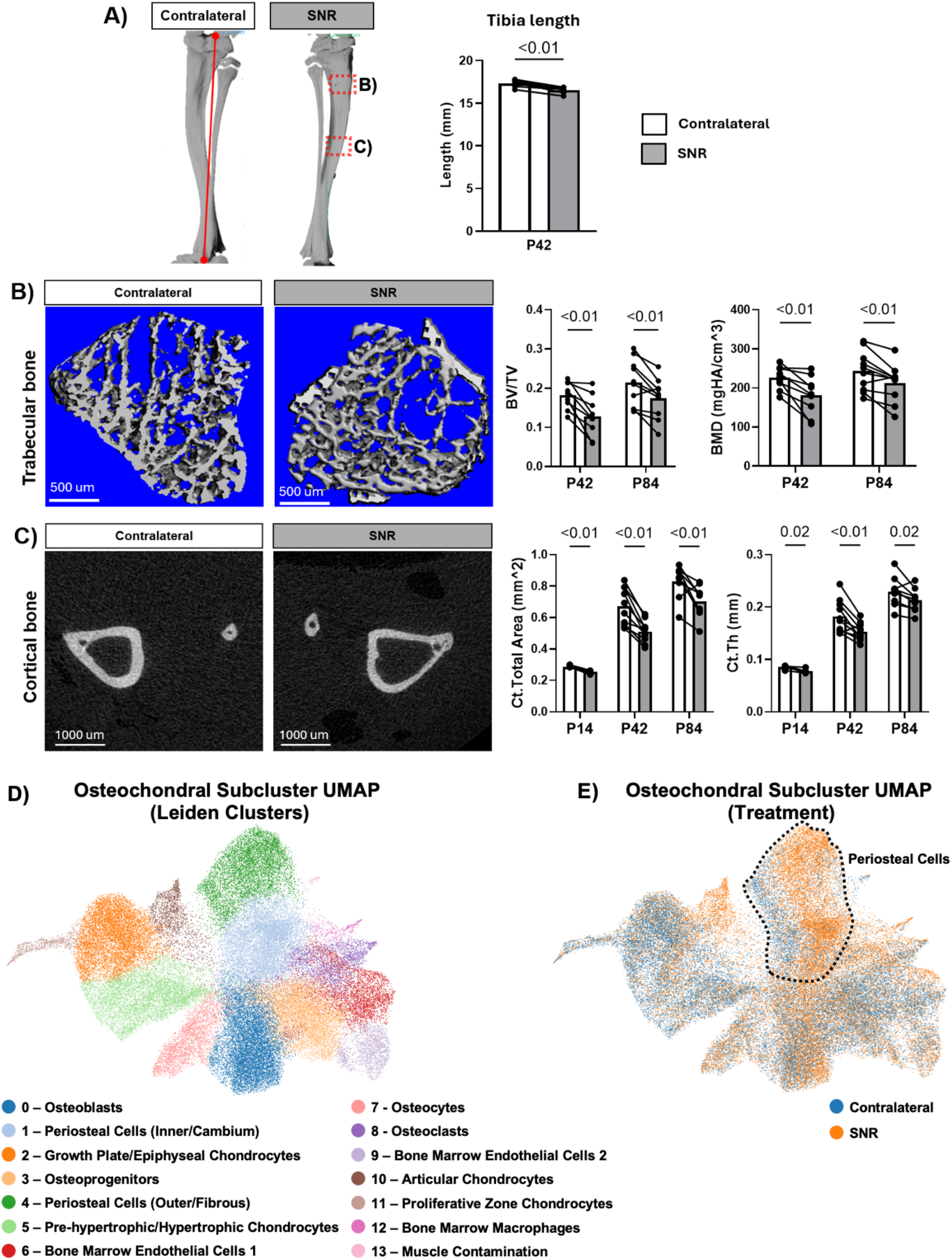
Reduced trabecular and cortical bone parameters with corresponding transcriptional changes in periosteal cells following SNR. A) SNR resulted in significantly shorter tibiae as measured from 3D reconstructed microCT images (red line) at postnatal day 42. B) SNR reduced trabecular bone parameters (BV/TV and BMD) at postnatal day 42 and P84. C) Cortical bone parameters (Cortical Total Area and Ct. thickness) were significantly reduced at P14, P42, and P84 in the SNR limb compared to the contralateral limb (n = 8-10, p<0.05). D) Reanalysis of osteochondral-associated cells identified 14 transcriptionally distinct subclusters. E) Osteochondral subcluster UMAP colored by treatment condition (SNR vs. contralateral), demonstrating a shift in periosteal cells (dashed line) (n = 3 mice/treatment).

We next sought to determine which cell populations in the bones were affected by the SNR procedure and could potentially explain the reduced bone growth. We reanalyzed the global chondrogenic (6) and osteogenic (3) clusters from the Xenium experiment, yielding 14 subclusters based on spatial location and expression of known markers (**Fig. 4D and S4**). Shifts in expression profiles in the periosteal (1 and 4) and articular chondrocytes (10) subclusters were most apparent. The higher number of articular chondrocytes in the SNR limb was due to SNR sections contained more of the talus articular cartilage (**Fig. S4A**). Specifically, within the osteogenic lineage, the expression patterns of both outer/fibrous and inner/cambium periosteal cells were most affected by the SNR procedure, with altered expression of genes related to mechano-signaling (*Acta2, Piezo2, Itga8, Emb,* and *Sfrp1*).

### SNR alters the growth dynamics and functional properties of the Achilles tendon

Next, we further dissected the effect of SNR on the overall growth of the Achilles tendon. Given the shift from proliferation- to matrix-driven growth in this early postnatal period (*25*, *27*), we first injected EdU intraperitoneally to assess cell proliferation at P4. EdU labeling revealed a reduction in proliferating cells in the SNR tendon compared to the contralateral limb (**Fig. 5A-B**, p < 0.01), suggesting that altered loading from SNR acutely suppresses cell proliferation-mediated growth at this early age. Next, we measured the cross-sectional area (CSA) of histological sections from the Achilles tendons up to P84 and found a smaller CSA in SNR compared to contralateral tendons at P14, P42, P56, and P84 (p < 0.001; **Fig. 5C-D)**. CSA increased over time in both groups; however, the overall growth trajectories were significantly lower in SNR tendons compared to contralateral controls **(Fig. 5D)**. Interestingly, the plantaris tendon CSA was similarly reduced in the SNR limb compared to the contralateral control **(Fig. S5)**, indicating that the altered loading environment affected positional as well as energy-storing tendons. The CSA reduction in the Achilles tendon was accompanied by a modest but significant increase in cell density at P14 (p = 0.01), P42 (p = 0.04), and P84 (p = 0.03) **(Fig. 5E)**. Given the reduction in cell proliferation at P4, this increase in cell density is likely indicative of reduced matrix accrual during these stages.

**Figure 5:**
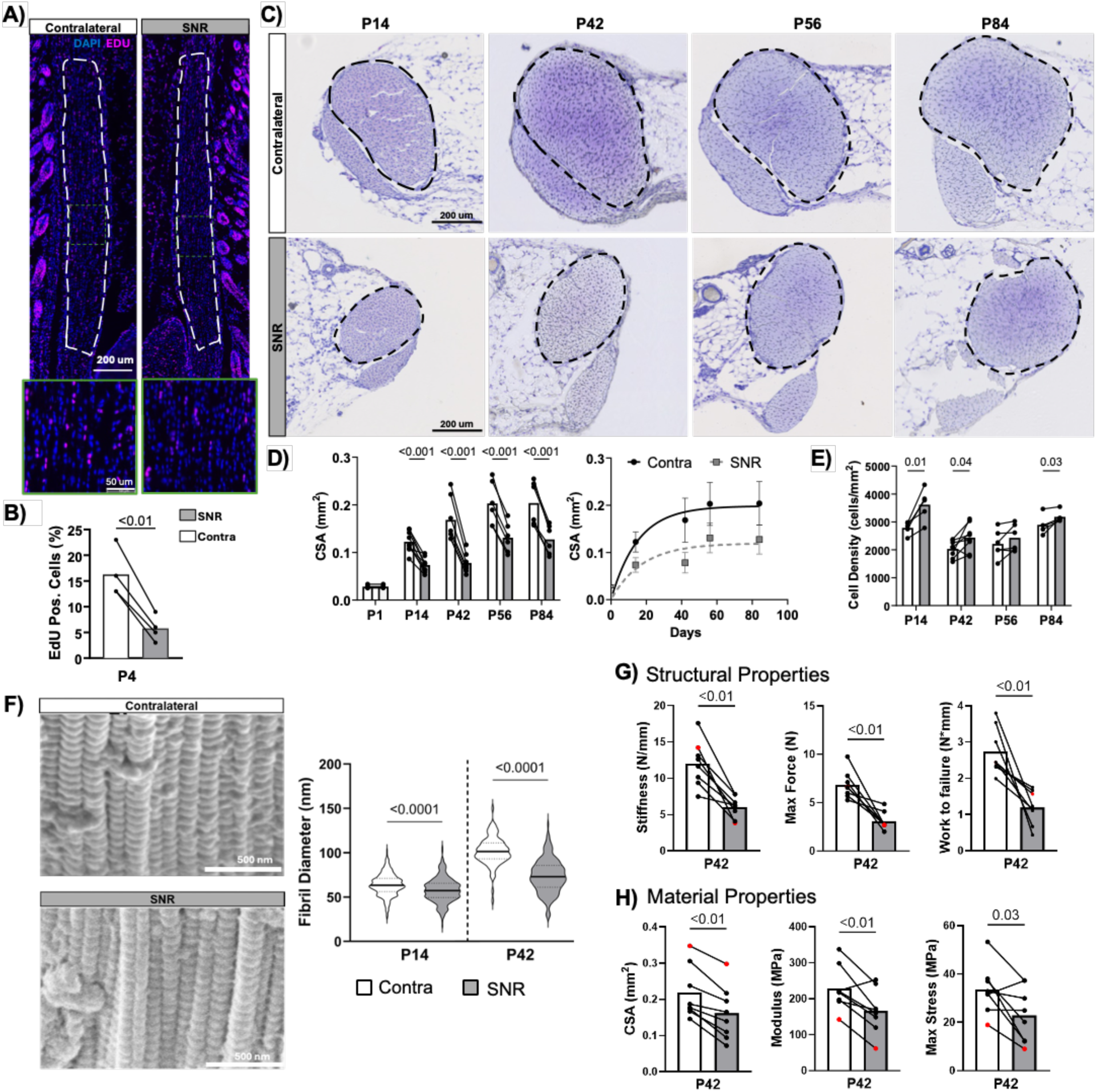
SNR impairs tendon growth, matrix assembly, and mechanical properties. A-B) EdU labeling revealed a reduction in proliferating cells in the SNR tendon compared to the contralateral limb (n = 4/group). C) Alterations in Achilles tendon morphometry following SNR shown by representative Toluidine Blue images. D) Reduced CSA in SNR tendons compared to contralateral at P14, P42, P56, and P84. Nonlinear regression with the von Bertalanffy growth function was used to model age-dependent changes in tendon cross-sectional area. Growth trajectories differed significantly between SNR and contralateral (contra) tendons (extra sum-of-squares F test, p<0.0001). E) Increased cell density in the SNR limb at P14, P42, and P84 (n = 6-9). F) Scanning electron microscopy (SEM) revealed a reduction in collagen fibril diameter following SNR (n = 3-4). G-H) Uniaxial tensile testing of the Achilles tendon demonstrated reduced structural properties (stiffness, max force, and work to failure), CSA, and material properties (modulus, max stress), following SNR (n=8, p < 0.05). Red dot indicates samples used for representative failure curves in Figure S7. Exact p-values are shown on the graph for statistically significant paired t-test comparisons.

To investigate matrix-driven growth, we assessed the expression of collagenous genes and the assembly of collagenous matrix. *Col1a1 (3.6 kb)-CFP* reporter mice were used to quantify *Col1a1* expression at single-cell resolution at P14, P42, P56, and P84 (**Fig. S6**). At P14, the SNR Achilles tendon displayed significantly reduced CFP intensity compared to the contralateral limb (p < 0.001), which is consistent with the reduced CSA at this time point (**Fig. 5C-D**). Interestingly, the SNR tendon displayed significantly higher CFP intensity compared to the contralateral control at P42 (p < 0.001), which remained elevated in the SNR tendon at P56 and P84. Overall, CFP intensities declined with age in both the contralateral and SNR limbs; however, the SNR limb showed a slower decline, retaining higher signal at later points (**Fig. S6A-B**). These general patterns also held true for endogenous *Col1a1* and *Col1a2* transcripts via bulk RNA sequencing (**Fig. S6C**). In addition to these transcriptional changes, the collagen fibril diameter distribution was significantly disrupted in the SNR tendons, as marked by reduced average diameter (**Fig. 5F**, p < 0.001) We next determined where these alterations in matrix production and assembly corresponded to changes in mechanical properties, as measured by uniaxial failure testing of the Achilles tendon. Both tissue-level structural (e.g., stiffness, max force) and material (e.g., modulus, max stress) properties were significantly reduced in the Achilles tendon of the SNR limb compared to the contralateral control (**Fig. 5G-H**, p < 0.05). Specifically, the SNR tendons exhibited a prolonged toe region, indicating delayed load transfer, as well as reduced stiffness, maximum force, and work to failure (p < 0.05). SNR also yielded tendons with reduced cross-sectional area in addition to reduced elastic modulus and maximum stress at P42 (p < 0.05).

These findings support the hypothesis that SNR limits matrix accrual and fibril thickening, which likely contributes to the impaired growth and the observed reduction in mechanical properties.

### SNR causes marked spatiotemporal changes in the expression profile of tendon cells

To define tendon-specific transcriptional changes at single-cell resolution, we restricted the Xenium dataset to a region of interest (ROI) encompassing the Achilles tendon, including the peritenon (**Fig. S8**). After quality filtering, a total of 40,000 tendon cells were analyzed, resulting in 10 Leiden clusters (**Fig. 6A**). Based on spatial localization, cluster composition, and canonical marker expression, clusters were annotated as: 4 different internal fibroblast (IF) populations, peritenon fibroblasts, perivascular cells, enthesis cells, macrophages, lymphatic cells, and myotendinous junction (MTJ) fibroblasts. Top-ranked marker genes for each Leiden cluster are shown in **Figure S8B**. The corresponding UMAP annotated for surgical treatment demonstrated a marked shift in the internal fibroblast populations (**Fig. 6B**). Among the IF subclusters, subcluster 0 corresponded to IF cells enriched in the SNR tendons, whereas subcluster 4 was enriched in the contralateral tendons, while subclusters 2 and 3 did not have a strong association with surgical treatment (**Fig. 6A-B**). IF4 (subcluster 4, contra enriched) was defined by higher expression of genes associated with metabolic and mitochondrial adaptation, including *Slc16a3, Higd1a*, and *Ndufa4l2*, along with proteoglycan and interface-associated extracellular matrix (ECM) genes such as *Acan*, *Clec3a*, and *Prg4* (**Fig. S8B**). In contrast, SNR-enriched internal fibroblasts (Cluster 0) showed increased expression of genes associated with matricellular signaling and collagen organization, including *Thbs4, Angptl7, and Tnmd* (**Fig. S8**).

**Figure 6:**
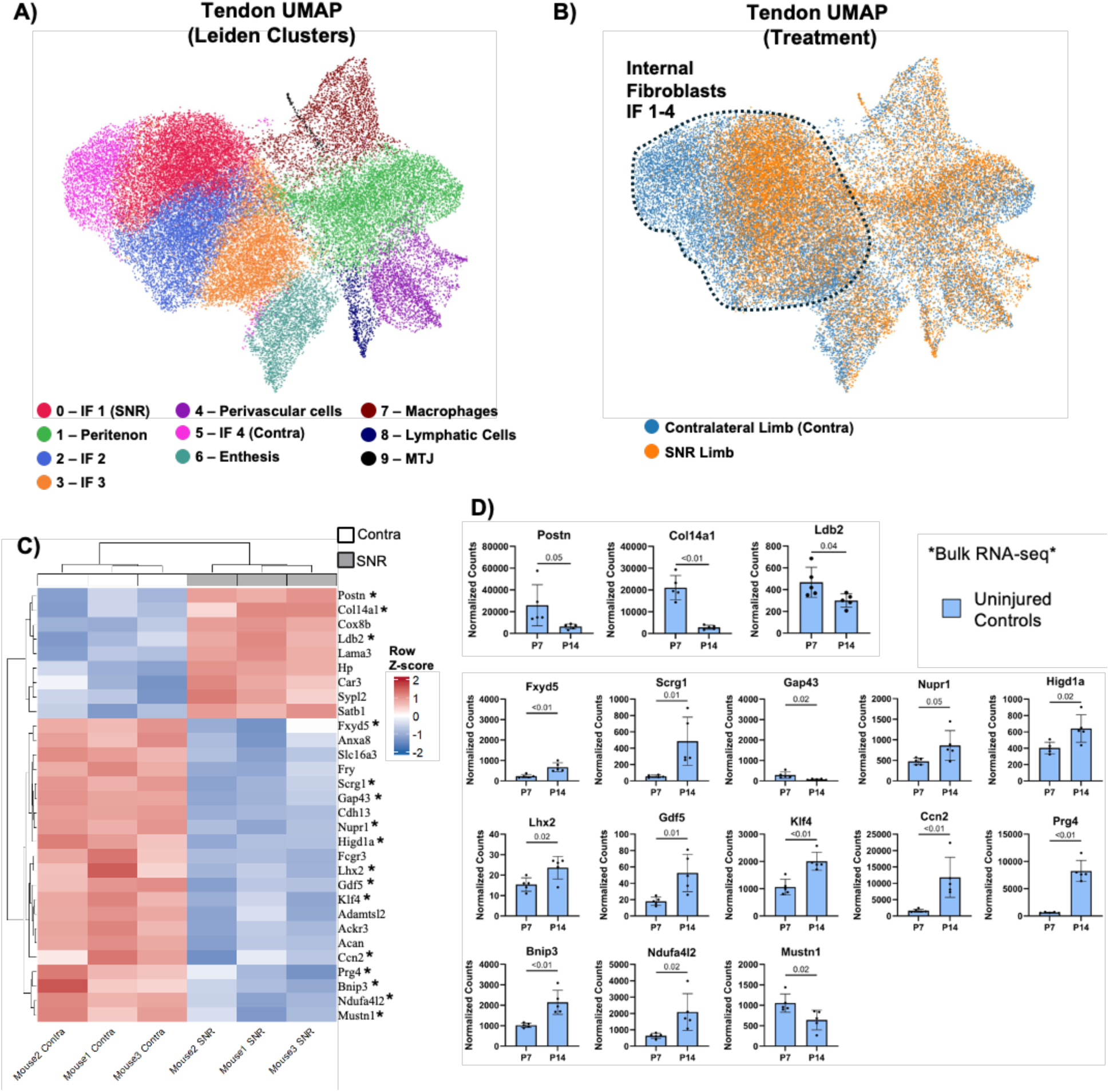
SNR alters expression profiles within the internal tendon fibroblasts. The Achilles tendon (AT) was segmented spatially to specifically analyze this tissue. A) UMAP of AT cells (see Figure S8) revealed 10 distinct Leiden clusters. B) UMAP of tendon subclusters colored by treatment condition. Dashed outline indicates IF clusters 1–4, which exhibited treatment-associated separation between contralateral and SNR tendons. C) Heatmap of differentially expressed genes between contralateral and SNR cells within the tendon IF populations (clusters 0, 2, and 4). Asterisks indicate genes that were also differentially regulated between P7 and P14 uninjured tendons in the bulk RNA-seq dataset. D) Bulk RNA-seq expression of the overlapping age-regulated genes in P7 and P14 control uninjured tendons (n = 5/group). Genes that normally increased from P7 to P14 were expressed at lower levels in SNR IFs, whereas genes that normally decreased with age were expressed at higher levels in SNR IFs.

Peritenon fibroblasts (cluster 1) displayed fewer transcriptional changes between SNR and contralateral limbs (**Fig. S8D**) with higher expression of *Postn, Ptn, Tnmd* in SNR cells and higher expression of *CCn3 Col8a1*, and *Vcan* in contralateral cells. Conversely, the enthesis cluster revealed minimal transcriptional differences with SNR treatment, although the sampling of this region was lower than the other clusters (**Fig. S8D**). These findings indicate that SNR has a cell-type-dependent effect, with the tendon internal fibroblast populations showing the most pronounced response.

To further probe the changes within the IF populations, we compared Xenium DEGs from tendon IF subclusters **(Fig. 6C)** to bulk RNA-seq data from control tendons at P7 and P14 **(Fig. 6D).** Of the 30 DEGs identified in the Xenium heatmap, 16 genes showed significant age-dependent changes in control tendons by bulk RNA-seq, with 5 genes decreasing and 11 genes increasing from P7 to P14. When compared to the Xenium dataset, SNR internal fibroblasts showed reduced expression of genes that normally increased from P7 to P14, and increased expression of genes that typically decreased with age in uninjured tendons (**Fig. 6C**, marked with *). Taken together, these findings indicate that SNR internal fibroblasts diverge from their normal developmental trajectory, with SNR tendons exhibiting shifts in internal fibroblast composition and gene expression consistent with a less mature tendon state (i.e., “younger” tendon) at P14.

The results thus far indicate an overall reduction in growth of the Achilles tendon with the internal tendon fibroblasts demonstrating delayed differentiation. To establish the global transcriptional changes of the tendon cells across several ages, we performed bulk RNA sequencing on control, contralateral, and SNR Achilles tendons collected at P7, P14, P42, and P56. Sequencing generated 40-150 million reads per sample. Principal component analysis (PCA) was performed using all expressed genes across all groups to identify major sources of transcriptional variance **(Fig. 7A)**. Age was the dominant driver of expression changes along PC1 (32.1% variance), with samples separating into distinct clusters in a temporal progression from P7 to P56. Notably, P42 and P56 samples clustered more closely together than at earlier time points, as expected given that growth rates were slowing between P42 and P56 (**Fig. 5D**). Surgical treatment correlated with expression changes along PC2 (13.3% variance), indicating a sustained effect of SNR on the tendon transcriptome. Along this axis, SNR samples formed distinct clusters from the age-matched contralateral and uninjured control tendons, which shared similar expression profiles (**Fig. 7A and S9C**). This finding is consistent with the bone parameters and tendon mechanics data, which show no significant differences between the control and contralateral limbs **(Fig. S10)**.

**Figure 7:**
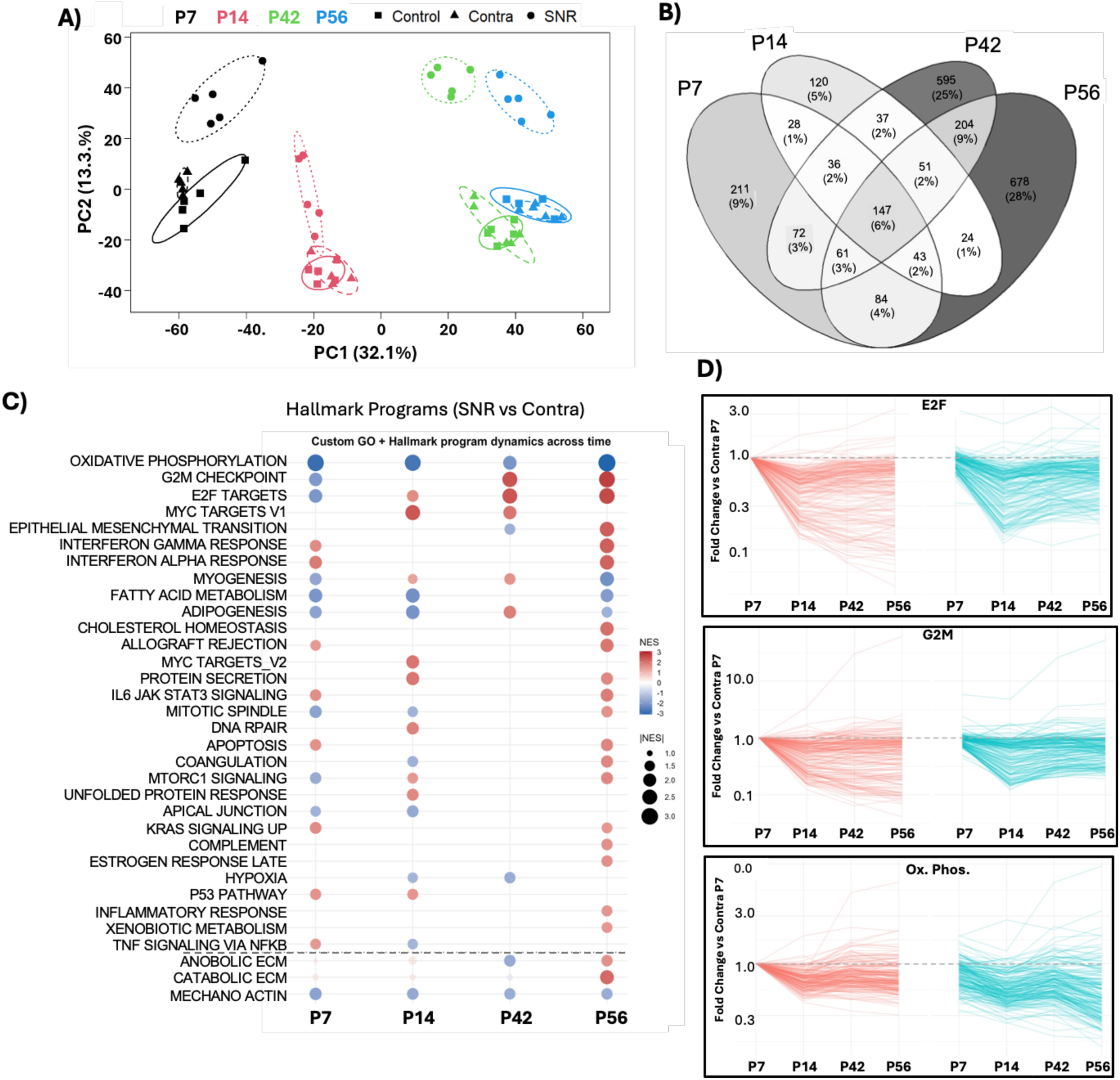
SNR produces distinct transcriptional signatures in the Achilles tendon during postnatal growth. A) Principal component analysis (PCA) of bulk RNA sequencing data, where SNR samples (circles) formed distinct clusters from the age-matched contralateral (triangles) and uninjured (squares) control tendons (n = 5 group). B) Venn diagram showing overlap of DEGs. C) GSEA of hallmark programs in SNR vs. contralateral tendons across time. Dot color indicates directionality of normalized enrichment score (NES) and dot size shows the magnitude of the NES. GSEA performed on tendon-curated custom GO programs is shown below the dashed line. D) Gene trajectory plots (normalized to the mean expression of the contralateral P7 sample) shows that the majority of cell cycle genes decreased with age in the contralateral AT but were mostly stable from P14 to P56 in SNR ATs. However, most genes in the oxidative phosphorylation (Ox Phos) hallmark were decreased at all time points, corresponding with the sustained downregulation in panel C.

We next sought to determine which gene sets were predominantly affected by SNR during postnatal growth. We identified differentially expressed genes (DEGs) between paired SNR and contralateral limbs within each time point. The majority of DEGs were unique to individual ages, with only 147 DEGs being common across all time points. Of these 147 genes, many were nerve- and muscle-specific, as expected given the SNR procedure and the likely inclusion of these cell types in the bulk samples. In addition, the number of DEGs increased with age (P7: 211, P14: 120, P42: 595, and P56: 678 DEGs), indicating a progressive phenotype in the SNR limbs. The greatest overlap was observed between P42 and P56, which shared 204 common DEGs (**Fig. 7B**). The overlap in DEGs between P42 and P56, combined with the closer PCA clustering of these later timepoints, suggests that transcriptional profiles converge during later stages of postnatal development. This pattern is consistent with the expected transition from early growth-associated programs toward a more mature tissue state.

Gene Set Enrichment Analysis (GSEA) comparing SNR and contralateral tendons at each time point demonstrated that Hallmark programs were most differentially regulated at P56. The top-enriched program was oxidative phosphorylation, which was markedly downregulated across all time points, indicating sustained alterations in metabolic gene programs following neonatal denervation. In addition, the cell cycle pathways, G2M checkpoint and E2F targets, were downregulated at P7, similar to our EdU results (**Fig. 5A-B**), but then increased at later time points **(Fig. 7C).** We used gene trajectory plots to determine whether pathway enrichment was driven by a small subset of highly regulated genes or whether it reflected coordinated shifts across most genes within the program. In this case, the majority of cell cycle genes in these pathways decreased with age in the contralateral tendons but, interestingly, remained stable from P14 to P56 in the SNR tendons (**Fig. 7D**). Given the lack of tendon-relevant hallmarks in GSEA, we performed GSEA on curated tendon-relevant programs encompassing anabolic and catabolic ECM remodeling, and mechanotransduction processes (**Table S1**). Anabolic ECM programs showed downregulation at P42 but upregulation at P56 in SNR tendons, which corresponded with the similar inflection point in the Col1a1-CFP reporter expression (**Fig. S6**). Catabolic ECM programs were also significantly upregulated in SNR at P56. In contrast, GSEA revealed stable suppression of mechanotransduction pathways in SNR tendons across all examined ages **(Fig. 7C)**. This persistent downregulation suggests reduced transduction of biophysical cues, likely due to impaired mechanical loading following the SNR procedure. Again, we found enrichment for myogenesis- and muscle contraction-related hallmark programs, likely reflecting skeletal muscle contamination in the bulk tendon samples. Interestingly, several inflammatory pathways (interferon alpha/gamma, IL6 JAK STAT, and inflammatory response) were all elevated in the SNR tendons at P56. These bulk RNA-sequencing results indicate profound dynamic changes in metabolic, mechanotransduction, and ECM-related pathways over time. The increased enrichment of anabolic, catabolic, and inflammatory programs at P56 also suggests that matrix remodeling may increase in SNR tendons as the phenotype progressively worsens.

## DISCUSSION

This study demonstrates that impaired neuromuscular control following neonatal sciatic nerve resection elicits dynamic, tissue-specific deficits in musculoskeletal tissue growth and maturation. Using SNR to reduce loading from early postnatal stages through maturity, we provide a multiscale analysis of the lower limb, including gait alterations, macro and micro-structure abnormalities, impairments in mechanical function, and transcriptional changes that correlate with altered growth trajectories. SNR produced distinct responses in muscle, bone, and tendon that together highlight the broad dependence of postnatal musculoskeletal development on mechanical cues. In muscle, SNR reduced muscle volume and shifted myogenic cells away from the normal mature fiber states and toward patterns consistent with fiber regeneration and tissue fibrosis. In bone, SNR impaired tibial and calcaneal growth, accompanied by modest transcriptional changes, primarily in the periosteum. SNR also resulted in pronounced alterations in tendon growth. Early after SNR, both cell proliferation and matrix synthesis were suppressed, and differentiation/maturation of the internal tendon fibroblasts was impaired. Interestingly, at later ages, some of these same processes (proliferation and matrix synthesis) were upregulated in SNR tendons relative to age-matched contralateral tendons. This altered developmental trajectory suggests that while SNR may reduce cumulative growth, it also causes a prolonged growth phase of the Achilles tendon. Together, these findings demonstrate that mechanical cues are required to sustain both the proliferative and matrix-accumulating phases of postnatal growth, which are required to establish the organized ECM necessary to withstand the demands of activities of daily living in adulthood.

Gross images revealed substantially reduced muscle growth following SNR, consistent with previous neonatal sciatic neurectomy studies showing that a loss of neural input during early postnatal development disrupts normal muscle maturation (*42*, *43*). Beyond reductions in size, we found altered transcriptional profiles of muscle cells in the Xenium dataset. SNR myogenic cells occupied transcriptional states distinct from those of the mature slow- and fast-twitch populations observed in the contralateral limb, consistent with disrupted muscle fiber maturation. In addition, the fibroblast and macrophage populations increased in SNR muscle, consistent with increased extracellular matrix remodeling and fibrosis. Histologically, increased centrally located nuclei and reduced phalloidin staining in the SNR muscle further supported ongoing muscle fiber remodeling.

While the importance of mechanical forces in bone growth and maintenance is well established, we analyzed osseous tissue growth in the distal limb to validate the sustained effects of reduced loading on osteogenesis. As expected, SNR had a negative impact on total bone growth (*33–35*), as demonstrated by reduced tibia length and trabecular and cortical bone parameters (p<0.05) in the SNR limbs. We detected deficits in cortical bone as early as P14 that persisted throughout the study (P84). We observed a decrease in tibial length and trabecular bone comparable to that reported in a recent study (*34*), and the reduction in trabecular parameters approaches the magnitude observed in more extreme mouse unloading models, such as tail suspension, which has been shown to produce 25–40% decreases in femoral and tibial BMD after 14 days of unloading (*44*, *45*). Interestingly, these substantial structural changes were accompanied by comparatively modest spatial transcriptional changes, with the clearest differences being genes related to mechanosignaling in the periosteum. Altered expression in periosteum is expected in this model (*46*); however, we did not detect appreciable changes in the osteocytes or growth plate chondrocytes, though this is likely attributable to poor sampling of the proximal tibial growth plate in our Xenium sections.

In contrast to muscle and bone, the contribution of mechanical loading to postnatal Achilles tendon growth is poorly defined, particularly during the early postnatal period when the tendon is rapidly expanding and establishing its mature structure. We investigated changes to both cell proliferation and matrix-driven growth. SNR yielded temporal changes in cell proliferation with suppression at early ages, as measured by both EdU labeling and expression of cell cycle genetic programs (i.e., G2M checkpoint and E2F targets) followed by an increase at later ages that was mostly due to a slower decline in these cell cycle programs in the SNR tendons with age. This pattern indicates that reduced mechanical loading not only suppresses early proliferation but also alters the normal age-dependent downregulation of cell cycle programs, possibly indicating a delayed growth trajectory or a transition to higher cell turnover, which will be a focus of future studies.

Impaired matrix accumulation was another major contributor to the reduction in cumulative tendon growth. The approximately 40% reduction we observed in CSA following SNR is consistent with impaired matrix deposition rather than loss of tendon cells, resulting in increased cellularity relative to tissue size. Tendons from SNR limbs were not only smaller in cross-sectional area but also demonstrated impaired material properties with smaller diameter collagen fibrils, indicating poorer matrix quality in addition to reduced matrix accrual. Similar reductions in mechanical properties have been reported following underloading studies in adult animals (*47*, *48*). At the ultrastructural scale, diminished mechanical loading following SNR likely disrupts collagen fibril lateral growth and maturation, contributing to defects in fibril formation and organization. These findings suggest that SNR not only delays early fibrillogenesis but also limits subsequent fibril thickening, a process that is known to be load-sensitive (*49–52*). These alterations were coupled with shifts in *Col1a1* gene expression, with suppressed reporter expression at early ages, followed again by an increase at later ages, relative to the contralateral tendon. In normal postnatal development, collagen expression declines after peaking within the first month of age as tendons transition toward matrix organization and maturation (*27*, *53–57*). Together, these findings indicate that reduced mechanical loading from SNR delays the normal developmental downregulation of collagen synthesis and simultaneously impairs effective matrix assembly and maturation, ultimately resulting in smaller, mechanically inferior tendons.

Together, the cell-cycle and matrix-synthesis data indicate a shared temporal shift in which early growth programs were suppressed, but later became elevated, compared to the contralateral tendon. Nonetheless, this prolonged growth-associated state was insufficient to compensate for the early deficits, resulting in impaired cumulative tendon growth and mechanical function.

The inability of this prolonged response to restore normal tendon structure may reflect a sustained disruption in cell mechanosensing. A substantial body of *in vitro* and *ex vivo* work demonstrates that cytoskeletal tension and actin organization are central regulators of tendon cell homeostasis in response to applied mechanical load (*57–61*), yet how these mechanotransductive pathways operate *in vivo* during early postnatal tendon development remains less clear. Our GSEA data begin to address this gap, revealing stable suppression of mechanotransduction and actin cytoskeleton gene programs in SNR tendons from P7 to P56, likely in the internal tendon fibroblast population, which appears to be the predominant responder to altered loading. This sustained suppression indicates that cells in SNR tendons exhibit reduced mechanotransduction output in response to reduced mechanical input. This is notable, given that prior work found that stress-deprived tendons undergo an MMP-driven catabolic response, and while loading can prevent this response, this protective effect only occurs when the actin cytoskeleton is intact (*60–62*). We did not notice this shift towards catabolism in our study, except that pro-inflammatory and catabolic markers were increased in the SNR tendons at P56, suggesting that this sustained reduction in loading may eventually shift the cells towards a catabolic state.

We used video gait analysis to better understand the altered loading environment contributing to these tissue phenotypes. SNR limbs exhibited sustained gait perturbations compared to contralateral limbs. Impaired muscle contraction in the hamstrings and distal muscle groups from SNR compromised the ability to elevate the hock and generate push-off into the swing phase, resulting in greater dorsiflexion of the SNR limb. The increased ankle dorsiflexion, combined with the increases in knee extension we previously reported (*33*), (*33*)demonstrates that strains of the Achilles tendon/muscle unit may be higher during gait in the SNR limb. However, we do not know whether the muscle or the tendon resists these strains, although other studies report elongation of the tendon that may be due to changes at the myotendinous junction (*63*).

Nonetheless, these gait impairments persisted to later time points in this longitudinal cohort of mice, suggesting the absence of nerve regeneration, sustained reductions in hindlimb loading, and altered force transmission to the Achilles tendon.

Collectively, our findings establish mechanical input as a critical regulator of postnatal musculoskeletal growth, dictating the dynamic growth trajectories of muscle, bone, and tendon. Neonatal SNR produced tissue-specific consequences across the lower limb, with muscle shifting toward persistent regenerative and fibrotic states, bone exhibiting substantial deficits in longitudinal and appositional growth, and the Achilles tendon following an altered growth trajectory characterized by reduced cumulative growth and a prolonged growth phase. This work also provides a comprehensive, multiscale dataset demonstrating how reduced loading alters tendon biology across length scales, from tissue structure to cellular behavior and transcriptional regulation. Ultimately, understanding the implications of altered loading on tissue growth and function could pave the way for targeted interventions to mitigate long-term deficits in musculoskeletal health from neurological disorders and impaired activity levels during this critical period of postnatal growth.

## MATERIALS AND METHODS

### Animal care and use

All animal housing, care, and procedures were approved by the University of Pennsylvania Institutional Animal Care and Use Committees (Protocol #806669). RNA-seq experiments were conducted with wildtype (WT) CD1 mice and 10x Xenium analyses were conducted in ScxCre;iMb2-Control-Mosaic mice (ScxCre: C57BL/6-Tg(Scx-GFP/cre)1Stzr/J; Mosaic: Gt(ROSA)26Sor^tm1(CAG-EYFP*,-mEYFP*,-tdTomato*,-mTFP1*)Ben^/J) mice (this line of mice is currently being used in a related unpublished study), while all other assays were performed in Col1a1(3.6kb)-CFP (Col1-CFP) reporter mice in a mixed CD1;CD57BL/6J background.

### Sciatic nerve resection

Unilateral sciatic nerve resection (SNR) was performed with microscopic guidance on postnatal day 1 (P1) mice, as previously reported (*33*, *36*). The sciatic nerve was accessed via a longitudinal incision through the skin on the posterior aspect of the femur. A segment of the nerve corresponding to ∼75% of the femoral length was removed to prevent regeneration of the nerve (*33*). Following nerve resection, the skin incision was closed with 6-0 suture.

### Gait analysis

Video gait analysis was performed using a custom-made walkway with a transparent bottom enclosed by 2 mirrors on either side to obtain video recordings from three viewpoints (ventral and left and right sagittal views). Ankle joint dorsiflexion angles were measured during two static frames of the gait cycle: paw strike and lift-off. Measurements made at paw strike used the first frame where the paw touched the ground (confirmed using both ventral and sagittal views). Measurements made at lift-off (hock height and ankle angle) used the last frame at which the paw was in contact with the walkway before entering the swing phase. Toe spread, defined as the distance between the first and fifth digits, was measured during the stance phase in the ventral view. Hock height was quantified using the sagittal view and was defined as the vertical distance from the ground (i.e., base plate) to the hock at the same time point. Longitudinal gait analysis was performed on the same cohort of male and female mice at postnatal days 14, 42, 56, and 84. *Tissue harvest:*

All mice were euthanized at designated time points via CO_2_ exposure according to the American Veterinary Medical Association’s guidelines for the Euthanasia of Animals. For micro-computed tomographic imaging and histological analyses, hindlimbs were fixed in 10% neutral buffered formalin for 2 days at 4℃, incubated in 30% sucrose for 1 day at 4℃, and imaged on the MicroCT and/or embedded in OCT and stored at -80℃. For biomechanical testing of the Achilles tendons, whole mice were stored at -20℃ following euthanasia until dissection on the day of testing.

### Multiplexed cryohistology

Cryosections (8 µm thick) of the OCT-embedded lower limb were created using Cryofilm 2C (Section-lab Co)(*64*) and stored at 4℃. Sections were fixed to glass slides using a Chitosan/UV glue adhesive. Achilles tendons were imaged with a Zeiss Axio Scan.Z1 slide scanner using a 10X objective. To assess fluorescent reporters, sections were counterstained with TO-PRO-3 (ThermoFisher, Cat#: T3605) and then imaged using Cy5 and CFP filters. Next, sections were stained with Toluidine Blue (0.025% solution, Sigma 364-M) and imaged under brightfield to assess overall structure at P1 (uninjured controls), P14, P42, and P56 (n=5-9/group). Cross-sectional area was obtained by drawing an outline around the Achilles tendon. Cellularity (# of nuclei per unit area) and fluorescent reporter (mean fluorescent intensity within each nucleus) quantification were determined from segmented nuclei (automated thresholding) using Fiji image analysis software. To assess cell proliferation, mice were given 3 consecutive daily injections of 5-ethynyl-2’-deoxyuridine (EdU) following surgery and sections were stained with Alexa Fluor 488 azide (Thermo Fisher Scientific) for EdU detection. Immunofluorescence at P42 was used to assess gastrocnemius muscle morphology using phalloidin, laminin, PDGFRα, and DAPI staining. Muscle and myofiber CSA, centrally nucleated fibers, and PDGFRα integrated fluorescence normalized to total myofiber area were quantified.

### Achilles tendon mechanical testing

Achilles tendons were dissected from the left SNR-treated hindlimb and the right contralateral limb (n*=*8 per group) at postnatal day 42. Sandpaper was placed 5 mm proximal to the calcaneal insertion and the calcaneus was placed in custom grips. Samples were loaded into a PBS bath on an Instron 5542 universal testing system. Mechanical testing was conducted in a saline bath where a preload to 0.005 N was applied followed by 10 cycles of preconditioning to 1.5% strain at 0.25 Hz, static stress relaxation (4% strain), and a ramp to failure at 0.05%/s.

### Micro-computed Tomographic Imaging

Excised hindlimbs were scanned on a Scanco μCT35 (voxel size=10.5 μm, energy=55 kV) scanner to measure tibial bone parameters at P14, P42, and P84 (n=8-10/group). Due to difficulty in measuring trabecular bone parameters reliably in P14 mice, trabecular bone parameters were quantified only at P42 and P84.

### Scanning Electron Microscopy (SEM)

Fixed frozen sections from the same mice used for cryohistology were fixed, dehydrated in a series of graded water-ethanol and ethanol-hexamethyldisilazane (HMDS) mixtures, and coated with a 6 nm thick platinum-palladium mixture and imaged via a Thermo Fisher Apreo 2S Lo Vac SEM to quantify collagen fibril diameter distributions (75-250 fibrils per mouse)(*65*).

### Bulk RNA Sequencing and Analysis

Achilles tendons were harvested from hindlimbs at P7, P14, P42, and P56 following sciatic nerve resection (SNR) or from contralateral control limbs. The Achilles tendon was carefully dissected using a stereomicroscope. Due to the small size, two Achilles tendons were pooled per biological replicate prior to RNA extraction. Immediately following dissection, fresh tendons were transferred directly into TRIzol reagent (Invitrogen) and homogenized using a mortar and pestle. Total RNA was isolated according to the manufacturer’s protocol. RNA concentration was measured using a Qubit fluorometer (Thermo Fisher Scientific). RNA amplification, ribosomal RNA (rRNA) depletion, library preparation, and sequencing were performed by Azenta/Genewiz (South Plainfield, NJ, USA). rRNA depletion was used to enrich for mRNA transcripts prior to library construction. Libraries were sequenced on an Illumina HiSeq platform to generate paired-end reads (2 × 150 bp). Sequencing reads were trimmed and aligned to the Mus musculus reference genome (GRCm38, Ensembl annotation). Gene-level read counts were generated from uniquely mapped reads (n=5 per group and time point).

Differential gene expression analysis was performed using DESeq2. Genes with an adjusted p-value < 0.05 and a log2 fold change greater than +1 or less than -1 were considered significantly differentially expressed genes. Gene Ontology (GO) analysis of these differentially expressed genes was performed using DAVID. Principal component analysis (PCA) and hierarchical clustering were performed to evaluate global transcriptional relationships among samples. Differential gene expression analysis between SNR and contralateral tendons was performed at each developmental time point.

### Gene Set Enrichment Analysis (GSEA)

GSEA was performed to identify coordinated changes in biological pathways for each timepoint. Genes were ranked using the Wald statistic from the DESeq2 analysis to define magnitude and variance. The ranked gene list was used as input for preranked GSEA analysis using Hallmark pathway gene sets from MSigDB to assess enrichment of a broad range of biological programs. Hallmark pathways were ranked by significance and normalized enrichment scores (NES) were calculated to reflect the degree to which genes from a given Hallmark were overrepresented for any given Hallmark. Hallmark pathways with an adjusted p-value < 0.05 were considered significantly enriched. In addition to Hallmark gene sets, custom gene programs relevant to ECM metabolism and mechanotransduction were defined using Gene Ontology (GO) annotations (gene list provided in **Table S1**. Specifically, collagen fibril organization, collagen biosynthetic process, and ECM assembly were used to define the Anabolic ECM program; while ECM disassembly, collagen catabolic process, ECM degradation, proteoglycan catabolic process, and catabolic process involved in ECM turnover were used to define the Catabolic ECM program.

The Mechano-Actin program was defined using GO Terms cellular response to mechanical stimulus, response to mechanical stimulus, mechanical stimulus, actin filament organization, actin cytoskeleton organization, stress fiber, actomyosin, and contractile fiber. We then used the same methods to determine significance and NES to observe how these gene sets changed over time.

### 10x Xenium Spatial Transcriptomics and Analysis

Cryofilm cryosections of Achilles tendons (n=3/group) at P14 were glued to 10x Xenium slides (two sections per mouse per slide, duplicated across two slides) (**Fig. S11**) with UV-activated adhesive (Norland Optical Adhesive 81), de-crosslinked with custom protease (ACD Bio) for 20 minutes at room temperature, stained using the 10x Genomics Cell Segmentation Add-on kit, and loaded into the Xenium Analyzer at the Children’s Hospital of Philadelphia single-cell core facility. Probes for 379 genes in the “Mouse Tissue Atlassing” panel, in addition to 100 custom genes (**Table S2**), were hybridized and imaged according to the 10x Xenium protocol.

Raw Xenium output was processed and analyzed using Scanpy and Squidpy. Cell-by-gene count matrices were loaded as .h5ad objects and analyzed using standard single-cell workflows. Counts were normalized and log-transformed prior to downstream analysis and all genes were retained for dimensionality reduction. Gene expression values were scaled prior to principal component analysis (PCA). PCA was performed on the full gene set, and the top 15 principal components were used to construct a k-nearest neighbor graph. Uniform Manifold Approximation and Projection (UMAP) embedding was computed from the neighborhood graph to visualize transcriptional relationships between cells. Unsupervised clustering was performed using the Leiden algorithm with a resolution parameter of 0.4 for the global UMAP (**Fig. 2A**), resolution of 0.2 was used for the muscle sub-clustering (**Fig. 3A**), resolution of 0.6 was used for the osteochondral sub-clustering (**Fig. 4D**), and resolution of 0.4 was used for the tendon-specific clustering (**Fig. 6A**), and cluster identities were assigned to individual cells. Spatial plots were generated from individual cells’ barcodes, and their associated metadata were used to identify treatment, Leiden cluster, mouse number, section, etc. Differential gene expression analysis was performed to identify cluster-specific marker genes. For each Leiden cluster, gene expression was compared against all remaining cells using pseudobulk analysis. Strong cluster-enriched marker genes were ranked using a scoring system incorporating both specificity and magnitude of expression. Gene specificity was defined as the difference between the proportion of cells expressing a gene within the target cluster (pct.1) and the proportion of cells expressing the gene in all other clusters (pct.2). This value was multiplied by the log2 fold change in expression between the target cluster and the remaining cells:

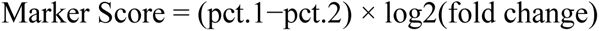

Cell density of specific muscle subclusters were measured by alpha-shape using the muscle cluster as the extent of the tissue range for each section and later grouped together by mouse and treatment resulting in n=3 per treatment. The alpha-shape area per section was used as the denominator and the number of cells occupying the defined area determined the cell density.

Marker gene heatmaps were generated in R using Seurat and ComplexHeatmap, based on raw transcript counts from cells assigned to each cluster and subcluster. Counts were log-normalized, and cluster-specific marker genes were identified using FindAllMarkers (Wilcoxon rank-sum test). For each cluster, the top 15 genes ranked by average log2foldchange were plotted, with expression values scaled using z-scores and ordered within each cluster to compare differences between Contra and SNR cells. Differential gene expression between Contra and SNR cells was assessed across all clusters and subclusters, with Contra cells as the reference group, and results were visualized as volcano plots, where upregulated genes are higher in SNR than in Contra, and downregulated genes are higher in Contra than in SNR.

### Statistics

SNR limb vs. contralateral limb groups were compared via paired t-tests at a=0.05. Shapiro-Wilk test of normal distribution was used to determine the extent of skewness, and transformation methods (e.g., logarithmic) were used to normalize the distribution of skewed variables. To assess the effects of two variables (surgery and development time points), we used a 2-way ANOVA with Tukey’s post hoc test for multiple comparisons when comparing three or more groups.

## ACKNOWLEDGEMENTS

This work was supported by NIH R01-AR075418, P50-AR080581, P50-AR080581-S1, and P30-AR069619 (histology, biomechanics, and MicroCT cores).

