## Supplemental Figures and Table 2 for "Impaired loading following denervation elicits dynamic, differential growth trajectories of musculoskeletal tissues"

A)

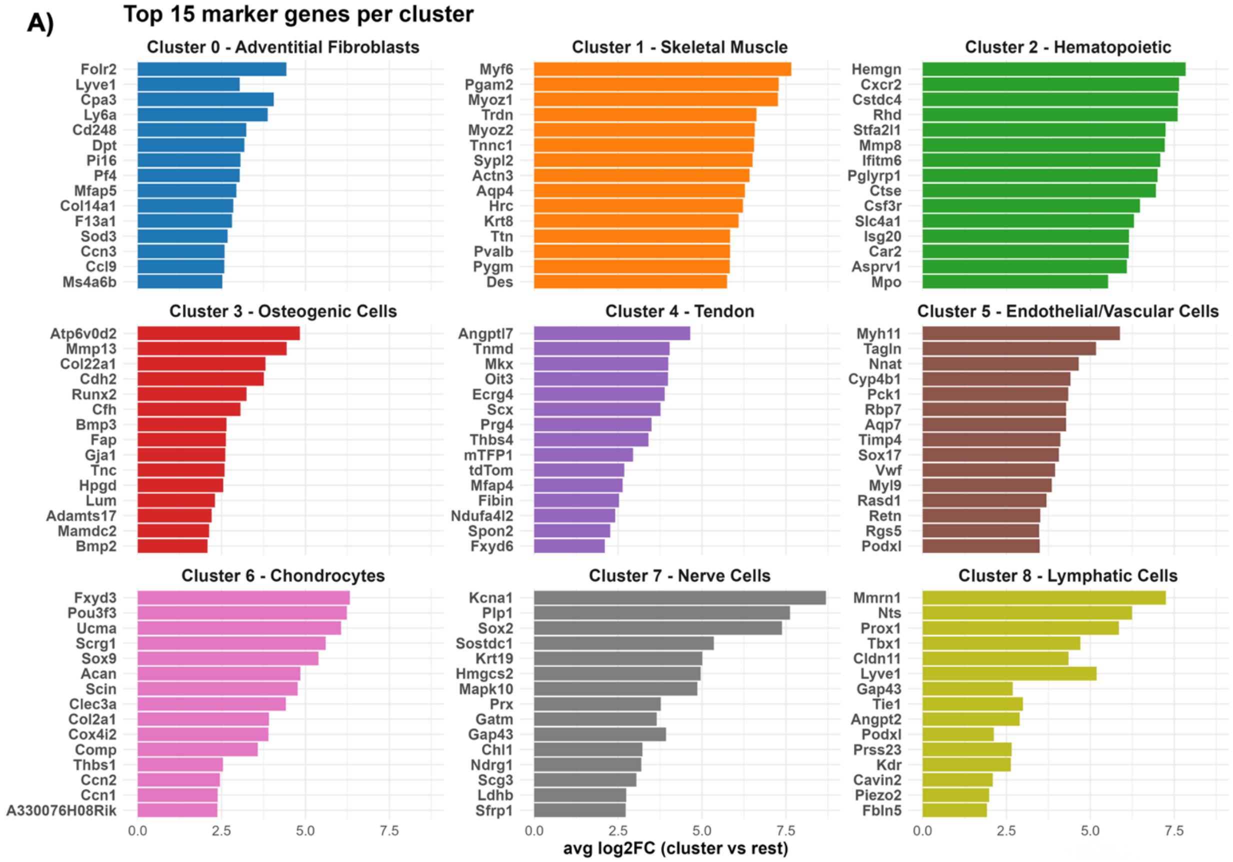

B)

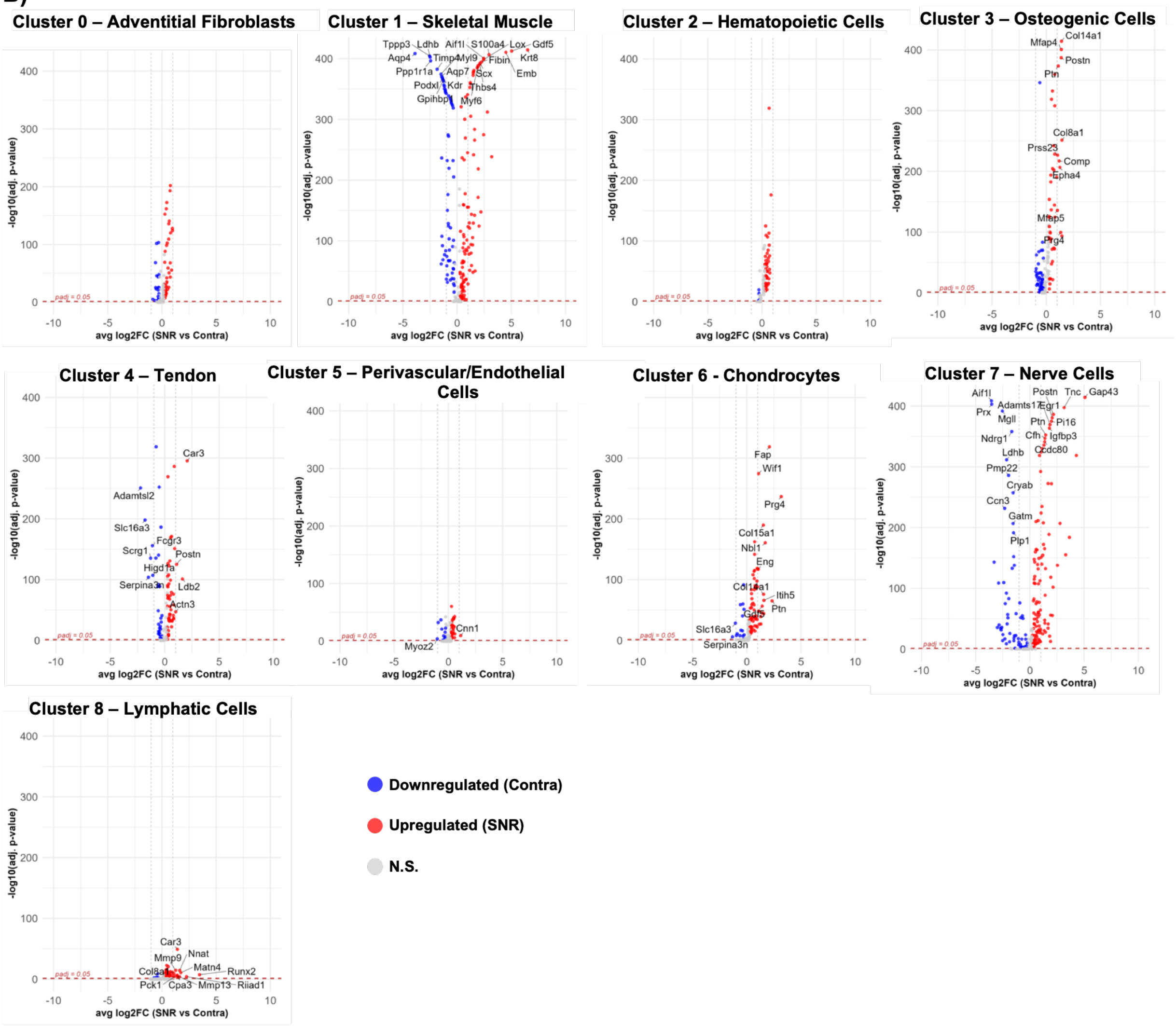

Cluster 0 – Adventitial Fibroblasts
Cluster 1 – Skeletal Muscle
Tppp3 Ldhb Aif1l S100a4 Lox Gdf5
Aqp4 Timp4 Myl9 Fibin Krt8
Ppp1r1a Aqp7 Scx Emb
Podxl Kdr Thbs4
Gpihbp Myf6
Cluster 2 – Hematopoietic Cells
Cluster 3 – Osteogenic Cells
Col14a1 Mfap4 Postn Ptn Col8a1 Prss23 Comp Epha4 Mfap5 Prg4
Cluster 4 – Tendon
Car3 Adamtsl2 Slc16a3 Fcgr3 Scrg1 Postn Higd1a Serpina3n Ldb2 Actn3
Cluster 5 – Perivascular/Endothelial Cells
Cnn1 Myoz2
Cluster 6 - Chondrocytes
Fap Wif1 Prg4 Col15a1 Nbl1 Eng Col1a1 Itih5 Ptn Gdf5 Slc16a3 Serpina3n
Cluster 7 – Nerve Cells
Aif1l Postn Tnc Gap43 Adamts17 Egr1 Prx Mgll Ptn Pi16 Ndrg1 Cfh Igfbp3 Ldhb Ccdc80 Pmp22 Cryab Ccn3 Gatm Plp1
Cluster 8 – Lymphatic Cells
Car3 Mmp9 Nnat Matn4 Col8a1 Runx2 Pck1 Cpa3 Mmp13 Riiad1
-log10(adj. p-value)
avg log2FC (SNR vs Contra)
padj = 0.05
Downregulated (Contra)
Upregulated (SNR)
N.S.

C)

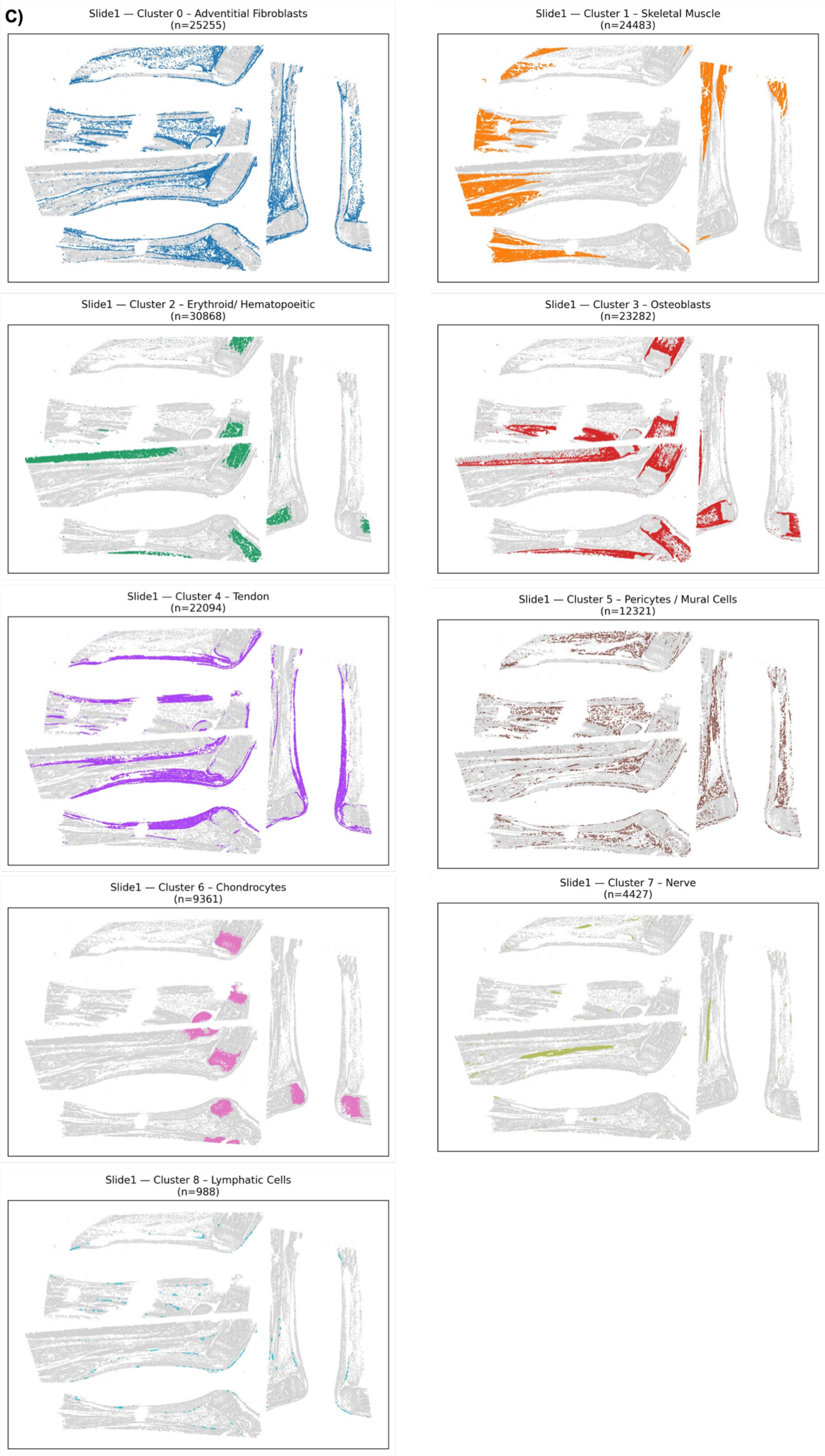

Slide1 — Cluster 0 – Adventitial Fibroblasts
(n=25255)
Slide1 — Cluster 1 – Skeletal Muscle
(n=24483)
Slide1 — Cluster 2 – Erythroid/ Hematopoeitic
(n=30868)
Slide1 — Cluster 3 – Osteoblasts
(n=23282)
Slide1 — Cluster 4 – Tendon
(n=22094)
Slide1 — Cluster 5 – Pericytes / Mural Cells
(n=12321)
Slide1 — Cluster 6 – Chondrocytes
(n=9361)
Slide1 — Cluster 7 – Nerve
(n=4427)
Slide1 — Cluster 8 – Lymphatic Cells
(n=988)

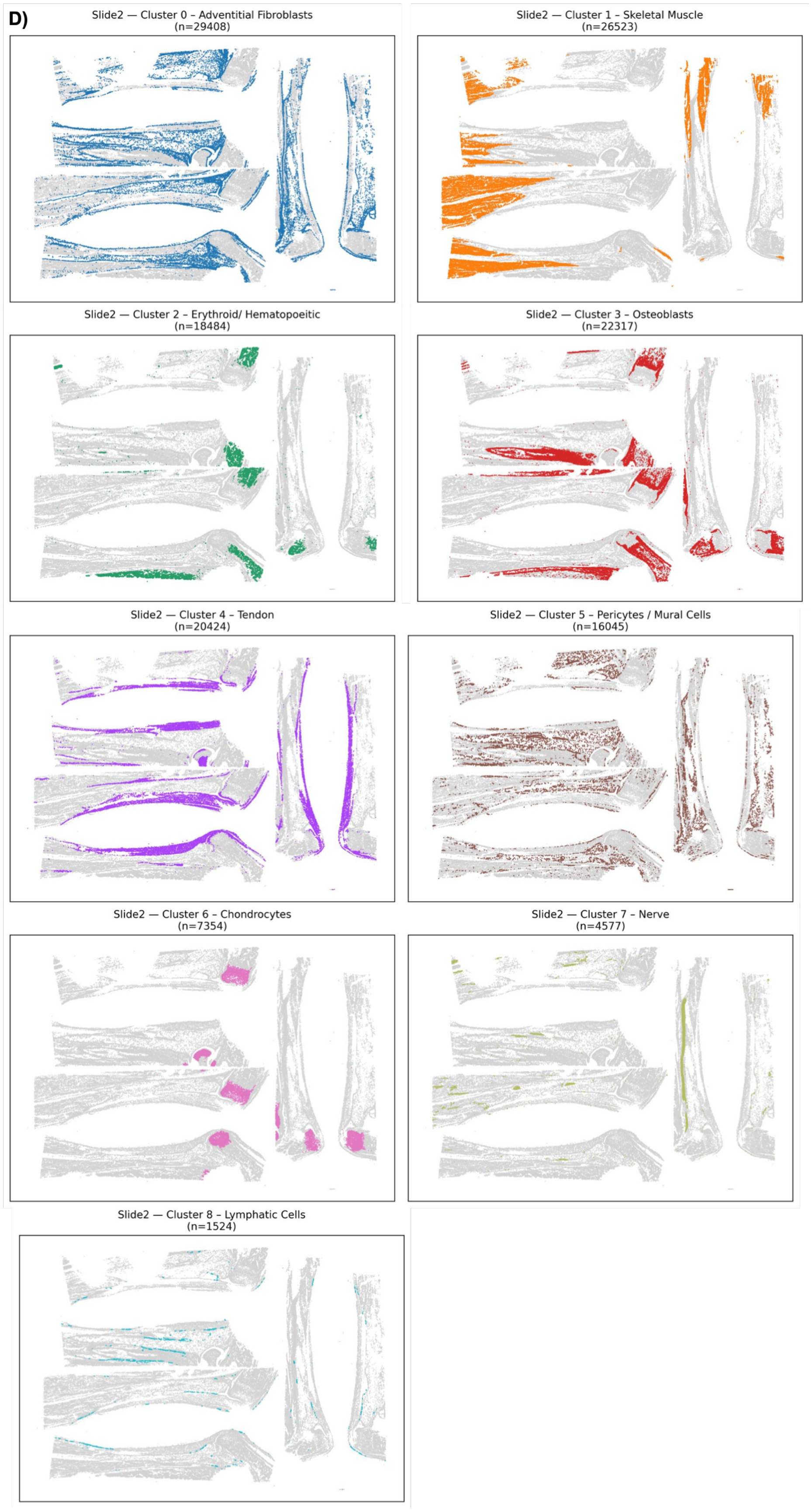
D)
Slide2 — Cluster 0 – Adventitial Fibroblasts
(n=29408)
Slide2 — Cluster 1 – Skeletal Muscle
(n=26523)
Slide2 — Cluster 2 – Erythroid/ Hematopoeitic
(n=18484)
Slide2 — Cluster 3 – Osteoblasts
(n=22317)
Slide2 — Cluster 4 – Tendon
(n=20424)
Slide2 — Cluster 5 – Pericytes / Mural Cells
(n=16045)
Slide2 — Cluster 6 – Chondrocytes
(n=7354)
Slide2 — Cluster 7 – Nerve
(n=4577)
Slide2 — Cluster 8 – Lymphatic Cells
(n=1524)

**Supplemental Figure 1: Global Xenium analysis of cell cluster identity, spatial distribution, and cluster abundance.** A) Top-ranked differentially expressed genes for each of the 9 Leiden clusters across tissue sections. B) Volcano plots showing differential gene expression between SNR and contralateral cells within each of the nine global Leiden clusters. Positive $\log_2$ fold-change values indicate genes enriched in SNR (red), whereas negative values indicate genes enriched in contralateral cells (blue) (n = 3). C) Spatial distribution of the nine Leiden clusters across all tissue sections from slide 1 and D) slide 2.

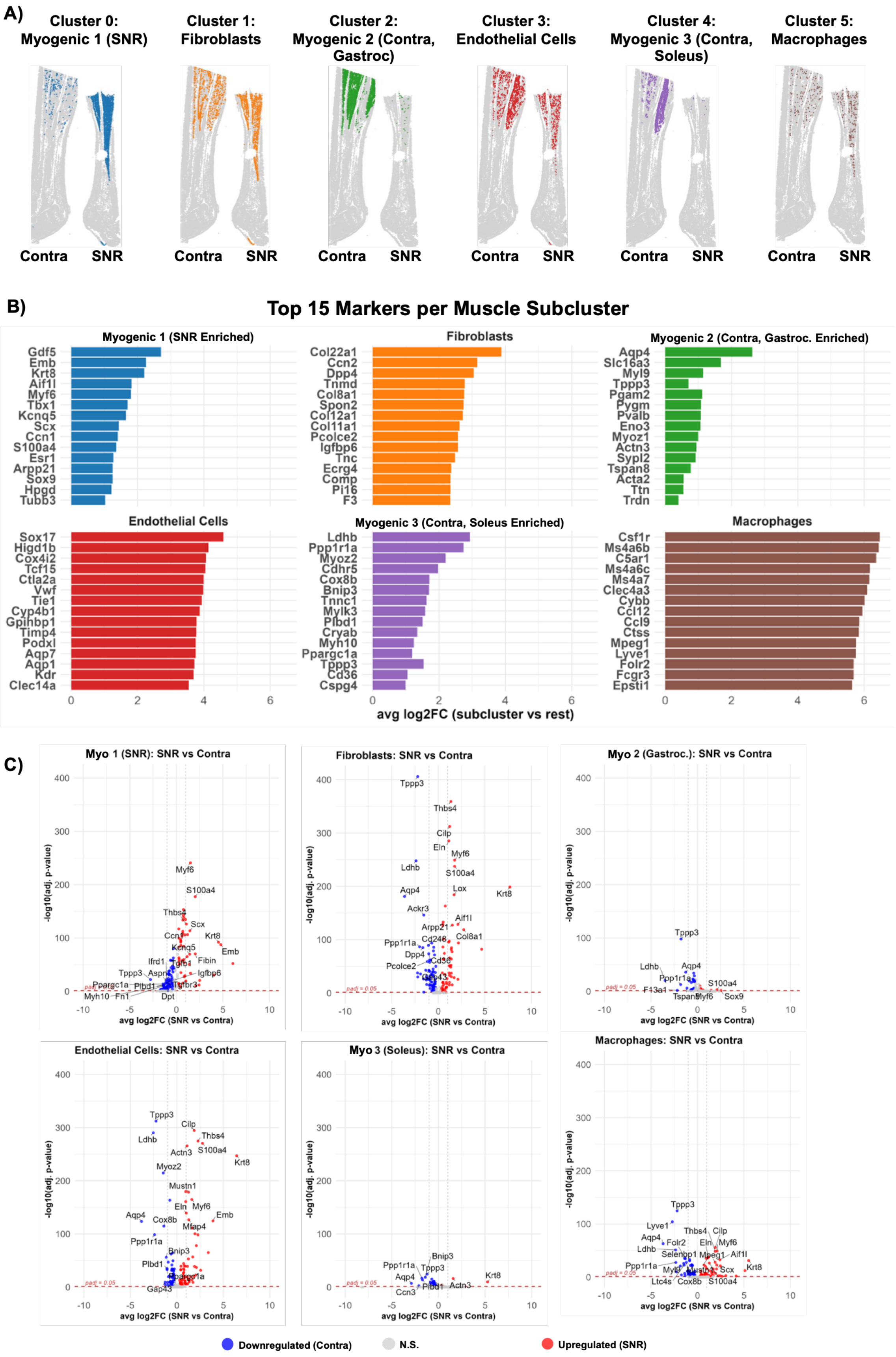
A)
Cluster 0: Myogenic 1 (SNR)
Cluster 1: Fibroblasts
Cluster 2: Myogenic 2 (Contra, Gastroc)
Cluster 3: Endothelial Cells
Cluster 4: Myogenic 3 (Contra, Soleus)
Cluster 5: Macrophages
Contra SNR
B)
Top 15 Markers per Muscle Subcluster
Myogenic 1 (SNR Enriched)
Gdf5 Emb Krt8 Aif1l Myf6 Tbx1 Kcnq5 Scx Ccn1 S100a4 Esr1 Arpp21 Sox9 Hpgd Tubb3
Fibroblasts
Col22a1 Ccn2 Dpp4 Tnmd Col8a1 Spon2 Col12a1 Col11a1 Pcolce2 Igfbp6 Tnc Ecrg4 Comp Pi16 F3
Myogenic 2 (Contra, Gastroc. Enriched)
Aqp4 Slc16a3 Myl9 Tppp3 Pgam2 Pygm Pvalb Eno3 Myoz1 Actn3 Sypl2 Tspan8 Acta2 Ttn Trdn
Endothelial Cells
Sox17 Higd1b Cox4i2 Tcf15 Ctla2a Vwf Tie1 Cyp4b1 Gpihbp1 Timp4 Podxl Aqp7 Aqp1 Kdr Clec14a
Myogenic 3 (Contra, Soleus Enriched)
Ldhb Ppp1r1a Myoz2 Cdhr5 Cox8b Bnip3 Tnnc1 Mylk3 Plbd1 Cryab Myh10 Ppargc1a Tppp3 Cd36 Cspg4
Macrophages
Csf1r Ms4a6b C5ar1 Ms4a6c Ms4a7 Clec4a3 Cybb Ccl12 Ccl9 Ctss Mpeg1 Lyve1 Folr2 Fcgr3 Epsti1
avg log2FC (subcluster vs rest)
C)
Myo 1 (SNR): SNR vs Contra
Fibroblasts: SNR vs Contra
Myo 2 (Gastroc.): SNR vs Contra
Endothelial Cells: SNR vs Contra
Myo 3 (Soleus): SNR vs Contra
Macrophages: SNR vs Contra
-log10(adj. p-value)
avg log2FC (SNR vs Contra)
padj = 0.05
Downregulated (Contra)
N.S.
Upregulated (SNR)

**Supplemental Figure 2: Skeletal muscle subcluster identity, spatial distribution, and treatment-associated gene expression following SNR**. A) Spatial mapping of the six skeletal muscle-associated subclusters, illustrating their anatomical distribution within the hindlimb. B) Top-ranked marker genes enriched in each skeletal muscle subcluster relative to the remaining subclusters. C) Volcano plots showing differential gene expression between SNR and contralateral cells within each of the six subclusters. Positive $\log_2$ fold-change values indicate genes enriched in SNR (red), whereas negative values indicate genes enriched in contralateral cells (blue) (n = 3).

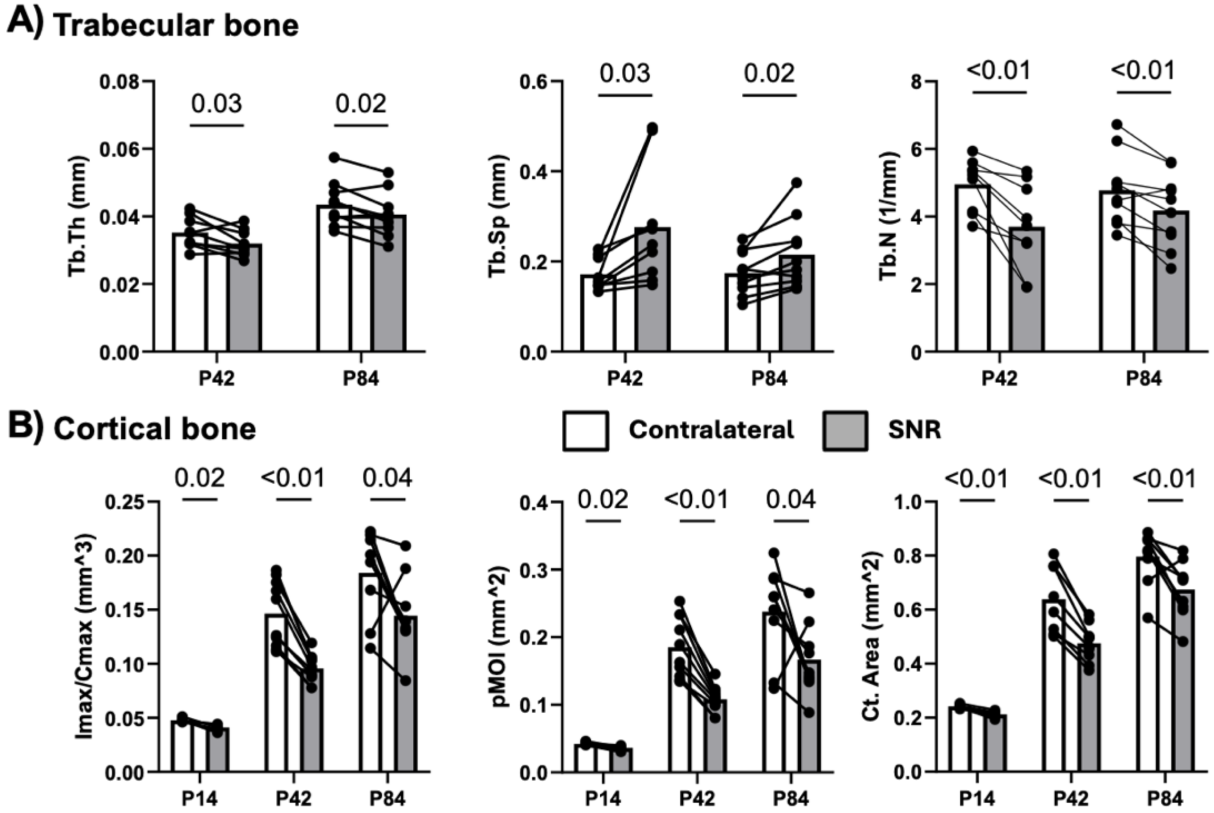

**Supplemental Figure 3: Neonatal SNR resulted in reduced bone parameters.** A) SNR yielded reduced trabecular bone parameters trabecular thickness (Tb.Th), spacing (Tb.Sp) and number (Tb.N) at postnatal day 42 and 84. B) Cortical bone parameters, including Imax/Cmax, polar moment of inertia (pMOI), and cortical area (Ct. area) were significantly reduced at P14, P42, and P84 (n=9-10, $p < 0.05$).

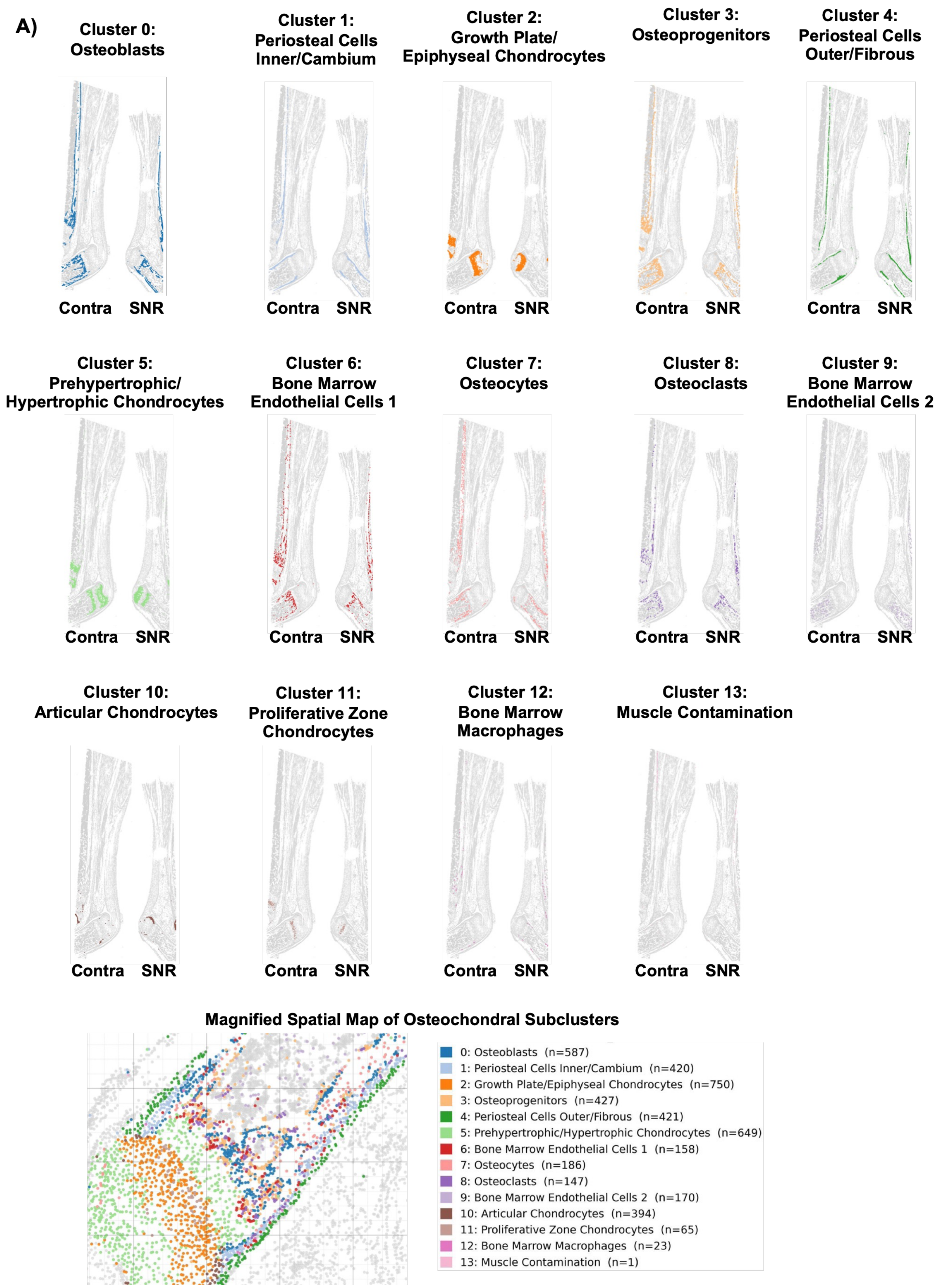

A)
Cluster 0: Osteoblasts
Cluster 1: Periosteal Cells Inner/Cambium
Cluster 2: Growth Plate/ Epiphyseal Chondrocytes
Cluster 3: Osteoprogenitors
Cluster 4: Periosteal Cells Outer/Fibrous
Cluster 5: Prehypertrophic/ Hypertrophic Chondrocytes
Cluster 6: Bone Marrow Endothelial Cells 1
Cluster 7: Osteocytes
Cluster 8: Osteoclasts
Cluster 9: Bone Marrow Endothelial Cells 2
Cluster 10: Articular Chondrocytes
Cluster 11: Proliferative Zone Chondrocytes
Cluster 12: Bone Marrow Macrophages
Cluster 13: Muscle Contamination
Contra
SNR
Magnified Spatial Map of Osteochondral Subclusters
0: Osteoblasts (n=587)
1: Periosteal Cells Inner/Cambium (n=420)
2: Growth Plate/Epiphyseal Chondrocytes (n=750)
3: Osteoprogenitors (n=427)
4: Periosteal Cells Outer/Fibrous (n=421)
5: Prehypertrophic/Hypertrophic Chondrocytes (n=649)
6: Bone Marrow Endothelial Cells 1 (n=158)
7: Osteocytes (n=186)
8: Osteoclasts (n=147)
9: Bone Marrow Endothelial Cells 2 (n=170)
10: Articular Chondrocytes (n=394)
11: Proliferative Zone Chondrocytes (n=65)
12: Bone Marrow Macrophages (n=23)
13: Muscle Contamination (n=1)

B)

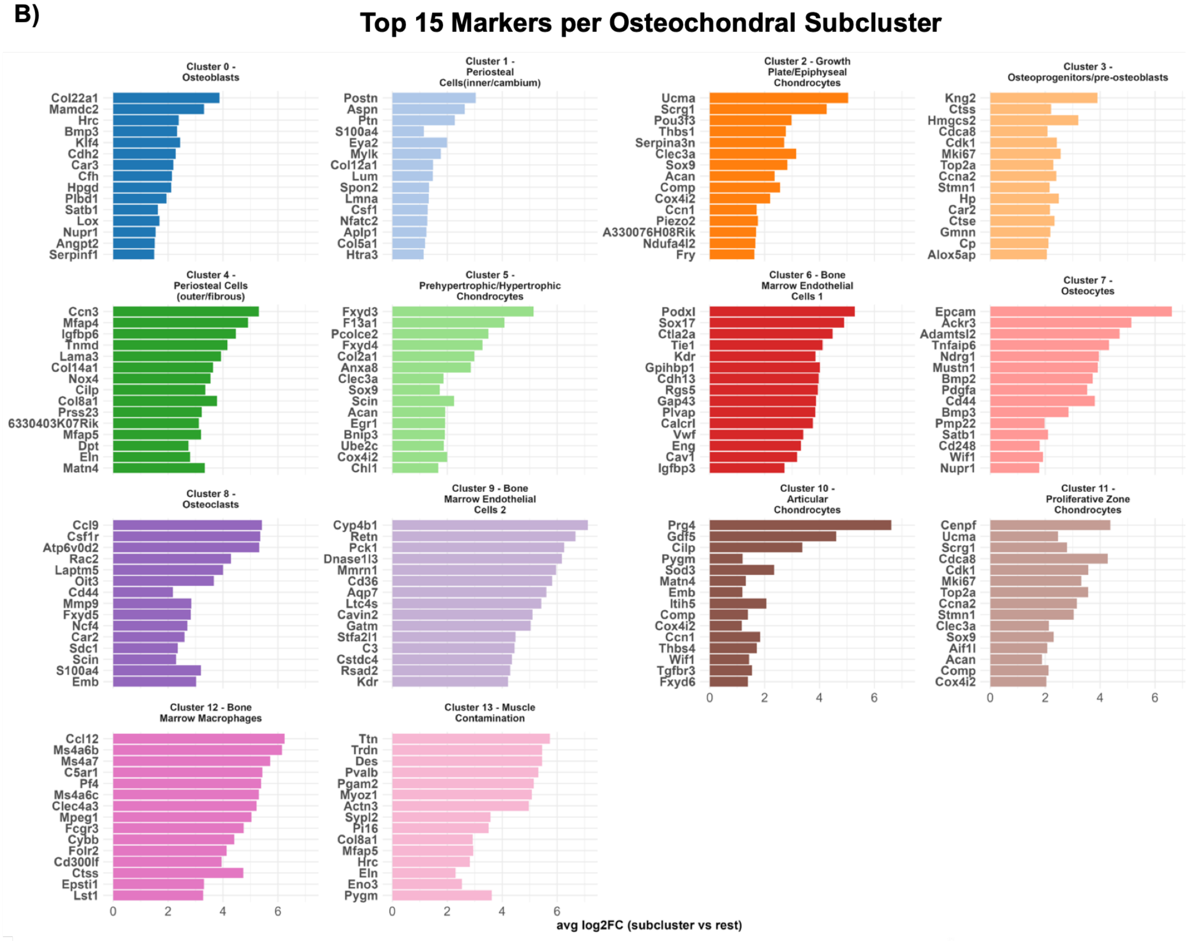

Top 15 Markers per Osteochondral Subcluster
Cluster 0 - Osteoblasts
Col22a1 Mamdc2 Hrc Bmp3 Klf4 Cdh2 Car3 Cfh Hpgd Plbd1 Satb1 Lox Nupr1 Angpt2 Serpinf1
Cluster 1 - Periosteal Cells(inner/cambium)
Postn Aspn Ptn S100a4 Eya2 Mylk Col12a1 Lum Spon2 Lmna Csf1 Nfatc2 Aplp1 Col5a1 Htra3
Cluster 2 - Growth Plate/Epiphyseal Chondrocytes
Ucma Scrg1 Pou3f3 Thbs1 Serpina3n Clec3a Sox9 Acan Comp Cox4i2 Ccn1 Piezo2 A330076H08Rik Ndufa4l2 Fry
Cluster 3 - Osteoprogenitors/pre-osteoblasts
Kng2 Ctss Hmgcs2 Cdca8 Cdk1 Mki67 Top2a Ccna2 Stmn1 Hp Car2 Ctse Gmnn Cp Alox5ap
Cluster 4 - Periosteal Cells (outer/fibrous)
Ccn3 Mfap4 Igfbp6 Tnmd Lama3 Col14a1 Nox4 Cilp Col8a1 Prss23 6330403K07Rik Mfap5 Dpt Eln Matn4
Cluster 5 - Prehypertrophic/Hypertrophic Chondrocytes
Fxyd3 F13a1 Pcolce2 Fxyd4 Col2a1 Anxa8 Clec3a Sox9 Scin Acan Egr1 Bnip3 Ube2c Cox4i2 Chl1
Cluster 6 - Bone Marrow Endothelial Cells 1
Podxl Sox17 Ctla2a Tie1 Kdr Gpihbp1 Cdh13 Rgs5 Gap43 Plvap Calcrl Vwf Eng Cav1 Igfbp3
Cluster 7 - Osteocytes
Epcam Ackr3 Adamtsl2 Tnfaip6 Ndrg1 Mustn1 Bmp2 Pdgfa Cd44 Bmp3 Pmp22 Satb1 Cd248 Wif1 Nupr1
Cluster 8 - Osteoclasts
Ccl9 Csf1r Atp6v0d2 Rac2 Laptm5 Oit3 Cd44 Mmp9 Fxyd5 Ncf4 Car2 Sdc1 Scin S100a4 Emb
Cluster 9 - Bone Marrow Endothelial Cells 2
Cyp4b1 Retn Pck1 Dnase1l3 Mmrn1 Cd36 Aqp7 Ltc4s Cavin2 Gatm Stfa2l1 C3 Cstdc4 Rsad2 Kdr
Cluster 10 - Articular Chondrocytes
Prg4 Gdf5 Cilp Pygm Sod3 Matn4 Emb Itih5 Comp Cox4i2 Ccn1 Thbs4 Wif1 Tgfbr3 Fxyd6
Cluster 11 - Proliferative Zone Chondrocytes
Cenpf Ucma Scrg1 Cdca8 Cdk1 Mki67 Top2a Ccna2 Stmn1 Clec3a Sox9 Aif1l Acan Comp Cox4i2
Cluster 12 - Bone Marrow Macrophages
Ccl12 Ms4a6b Ms4a7 C5ar1 Pf4 Ms4a6c Clec4a3 Mpeg1 Fcgr3 Cybb Folr2 Cd300lf Ctss Epsti1 Lst1
Cluster 13 - Muscle Contamination
Ttn Trdn Des Pvalb Pgam2 Myoz1 Actn3 Sypl2 Pi16 Col8a1 Mfap5 Hrc Eln Eno3 Pygm
0 2 4 6
avg log2FC (subcluster vs rest)

C)

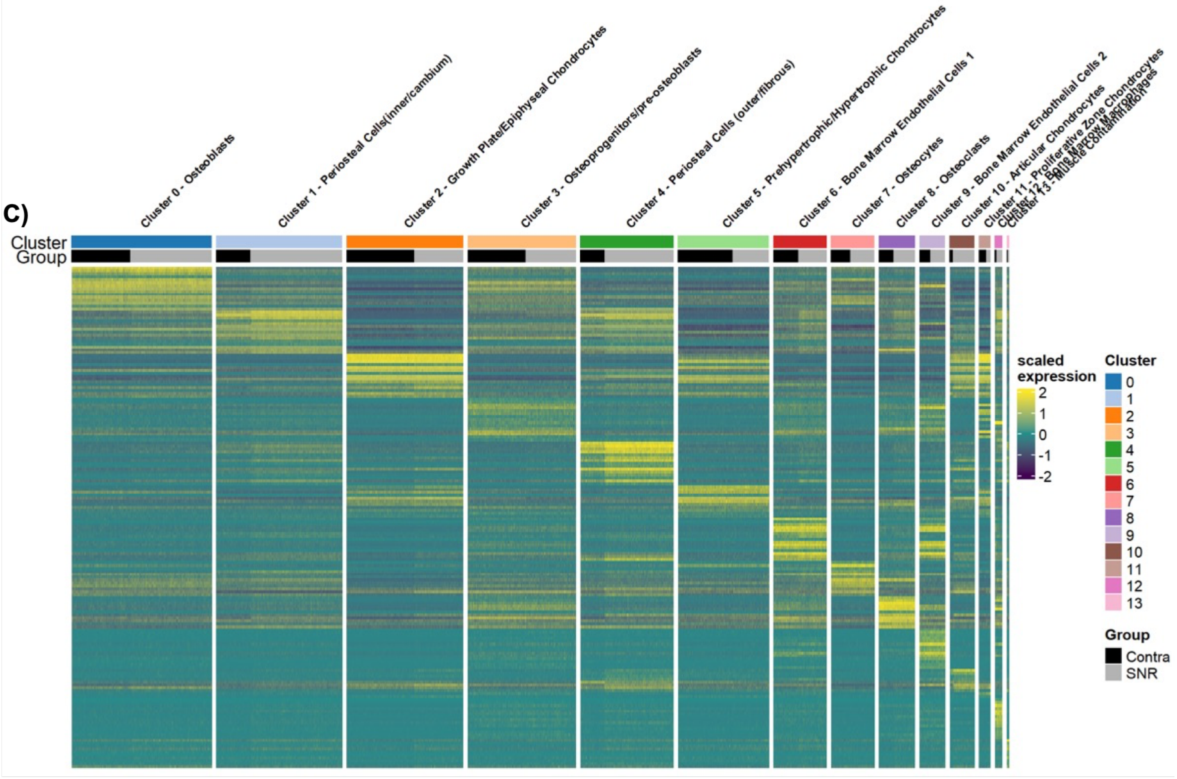

Cluster 0 - Osteoblasts
Cluster 1 - Periosteal Cells(inner/cambium)
Cluster 2 - Growth Plate/Epiphyseal Chondrocytes
Cluster 3 - Osteoprogenitors/pre-osteoblasts
Cluster 4 - Periosteal Cells (outer/fibrous)
Cluster 5 - Prehypertrophic/Hypertrophic Chondrocytes
Cluster 6 - Bone Marrow Endothelial Cells 1
Cluster 7 - Osteocytes
Cluster 8 - Osteoclasts
Cluster 9 - Bone Marrow Endothelial Cells 2
Cluster 10 - Articular Chondrocytes
Cluster
Group
scaled expression
2
1
0
-1
-2
Cluster
0 1 2 3 4 5 6 7 8 9 10 11 12 13
Group
Contra
SNR

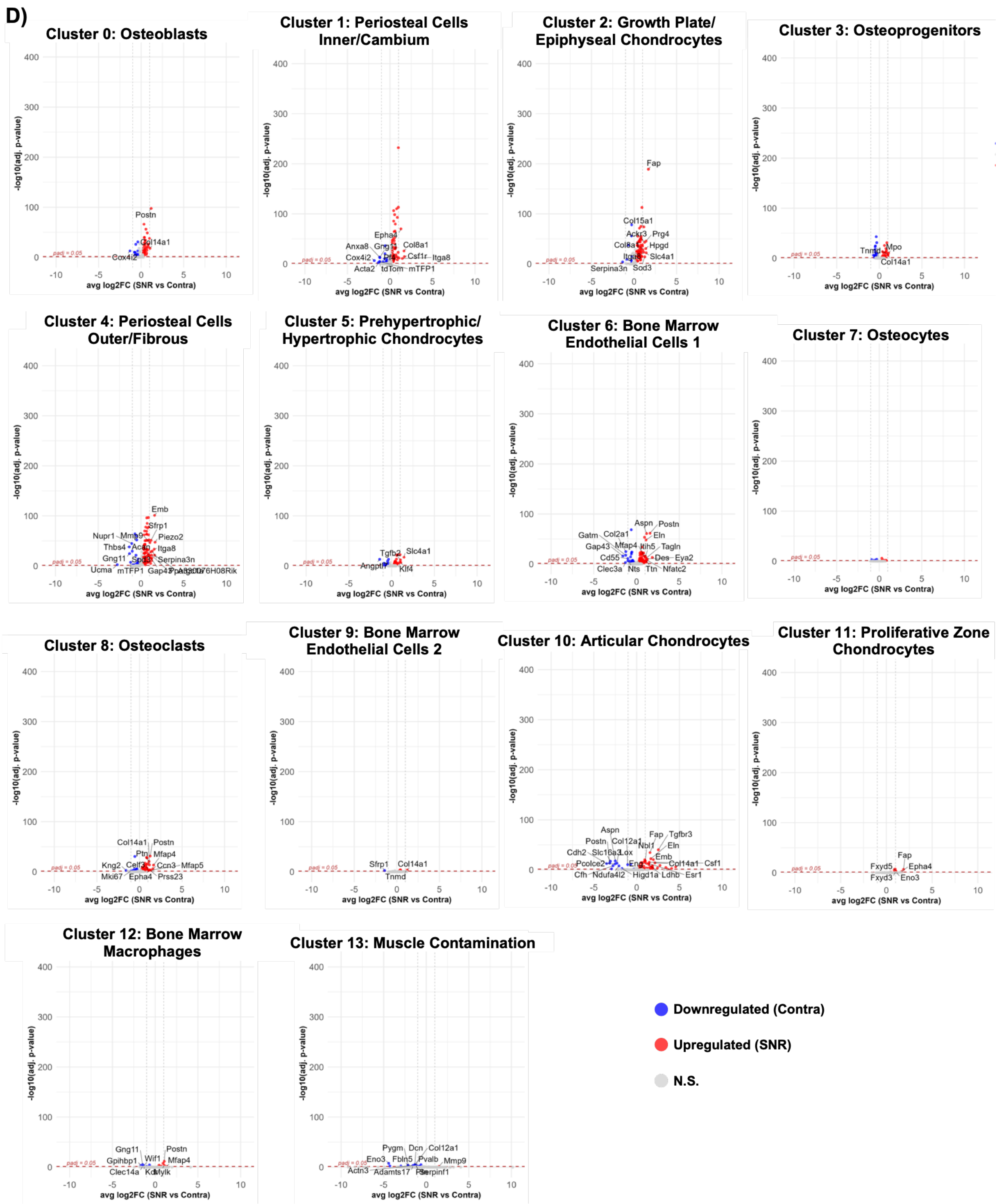

**Supplemental Figure 4: Osteochondral subcluster identity, spatial distribution, and treatment-associated gene expression following SNR.** A) Spatial mapping of the 14 osteochondral subclusters across tissue sections, illustrating their anatomical distribution within the bone and osteochondral regions. B) Top-ranked marker genes enriched in each osteochondral subcluster relative to the remaining subclusters. C) Heatmap of scaled gene expression across osteochondral subclusters and treatments, demonstrating cluster-specific expression patterns. D)

Volcano plots showing differential gene expression between SNR and contralateral cells within each of the 14 subclusters. Positive $log_2$ fold-change values indicate genes enriched in SNR (red), whereas negative values indicate genes enriched in contralateral cells (blue) (n = 3).

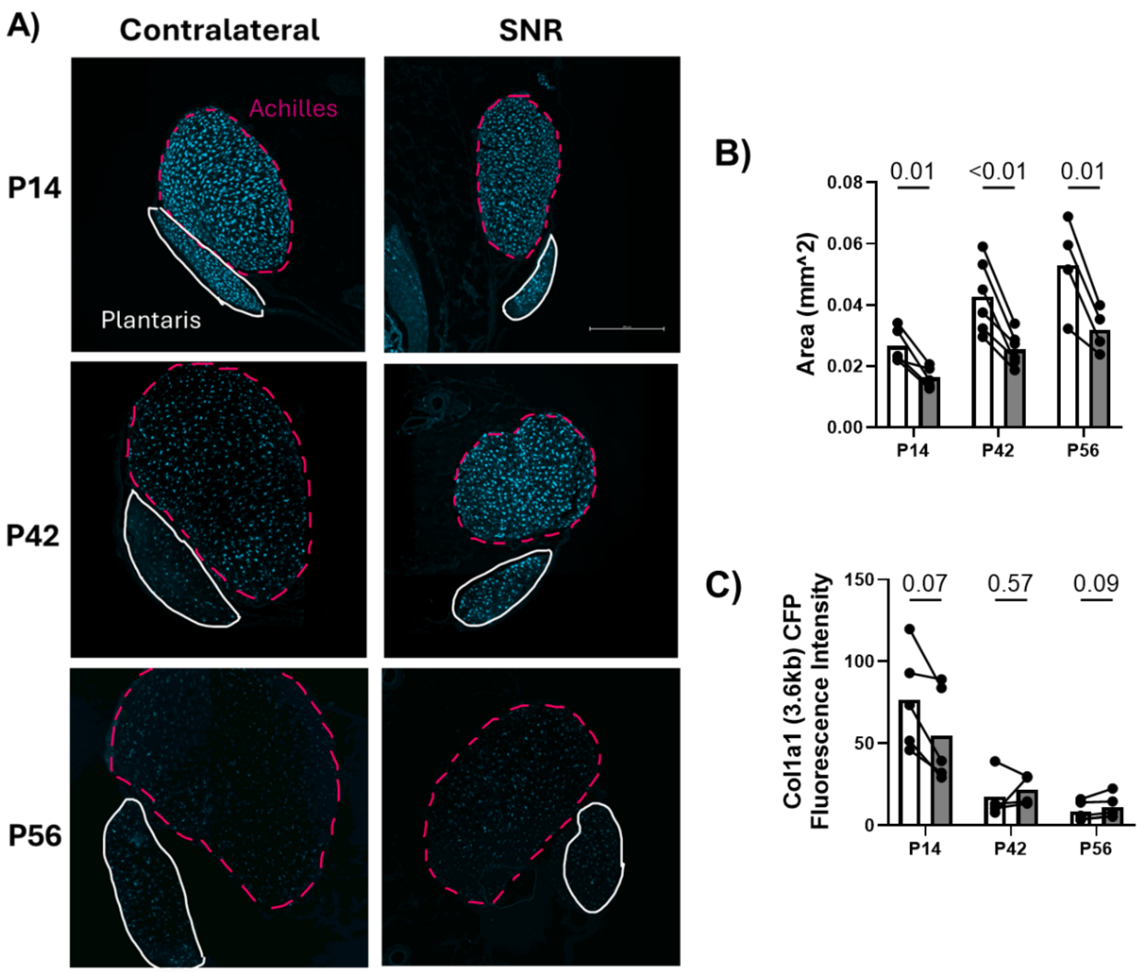

**Supplemental Figure 5: SNR alters the growth and maturation of the plantaris tendon.** A) Representative images of the Achilles (pink dashed line) and plantaris (white outline) tendons reveal B) significantly decreased cross-sectional area of the plantaris at all timepoints. C) *Col1a1(3.6kb)-CFP* expression was not significantly altered in the SNR limb compared to the contralateral limb (n=4-6, $p<0.05$).

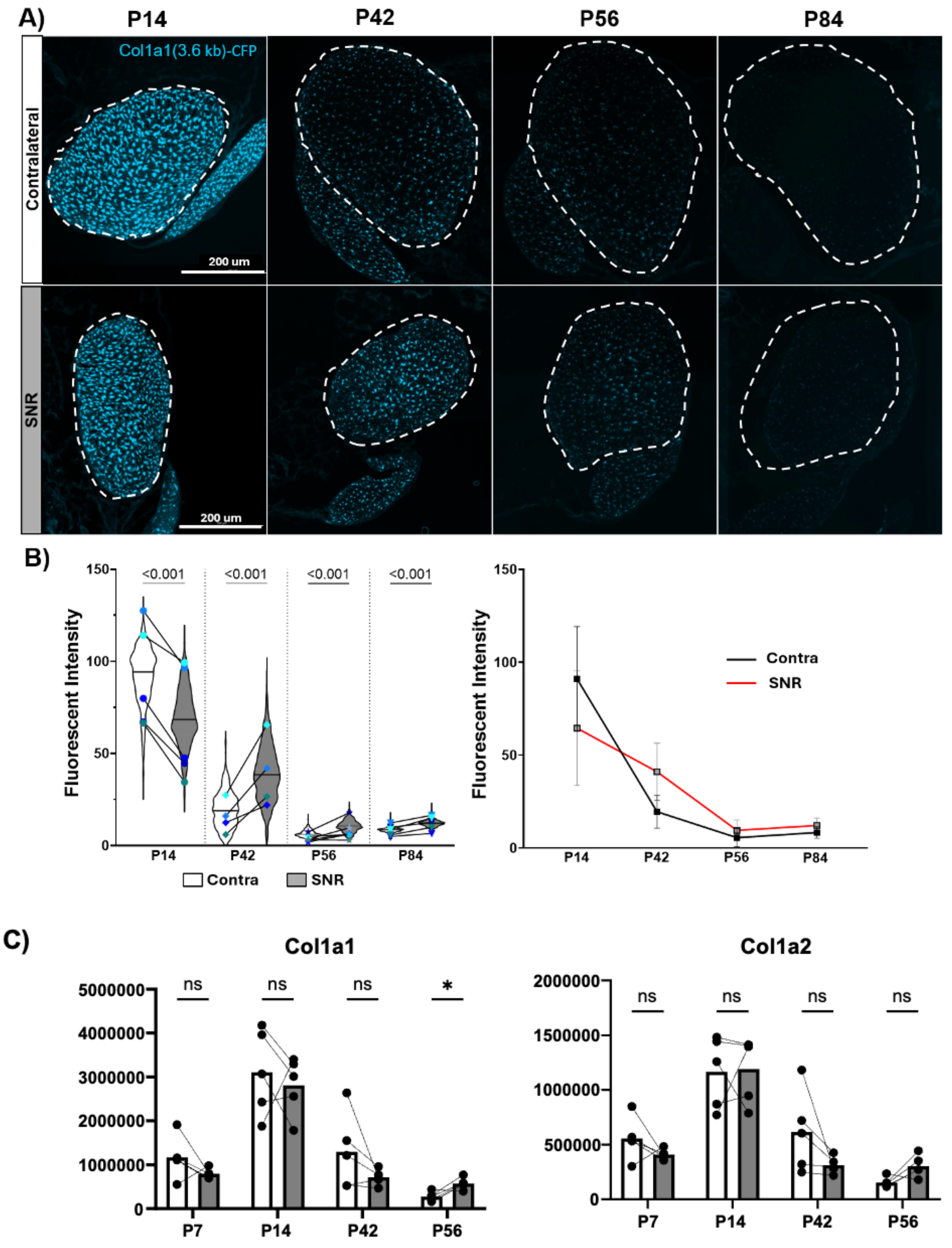

**Supplemental Figure 6: The temporal expression patterns of Col1a1(3.6kb)-CFP expression in tendon fibroblasts are altered following SNR.** A) SNR delays the temporal decline in (Col1a1 3.6kb) expression. B) Col1a1(3.6kb)-CFP expression was reduced in SNR tendons at P14 however the SNR limb exhibited significantly higher CFP intensity at later time points (P42, P56, and P84) compared to the contralateral limb (n=4-6). C) Expression of collagen genes from bulk RNA-seq across postnatal growth following SNR.

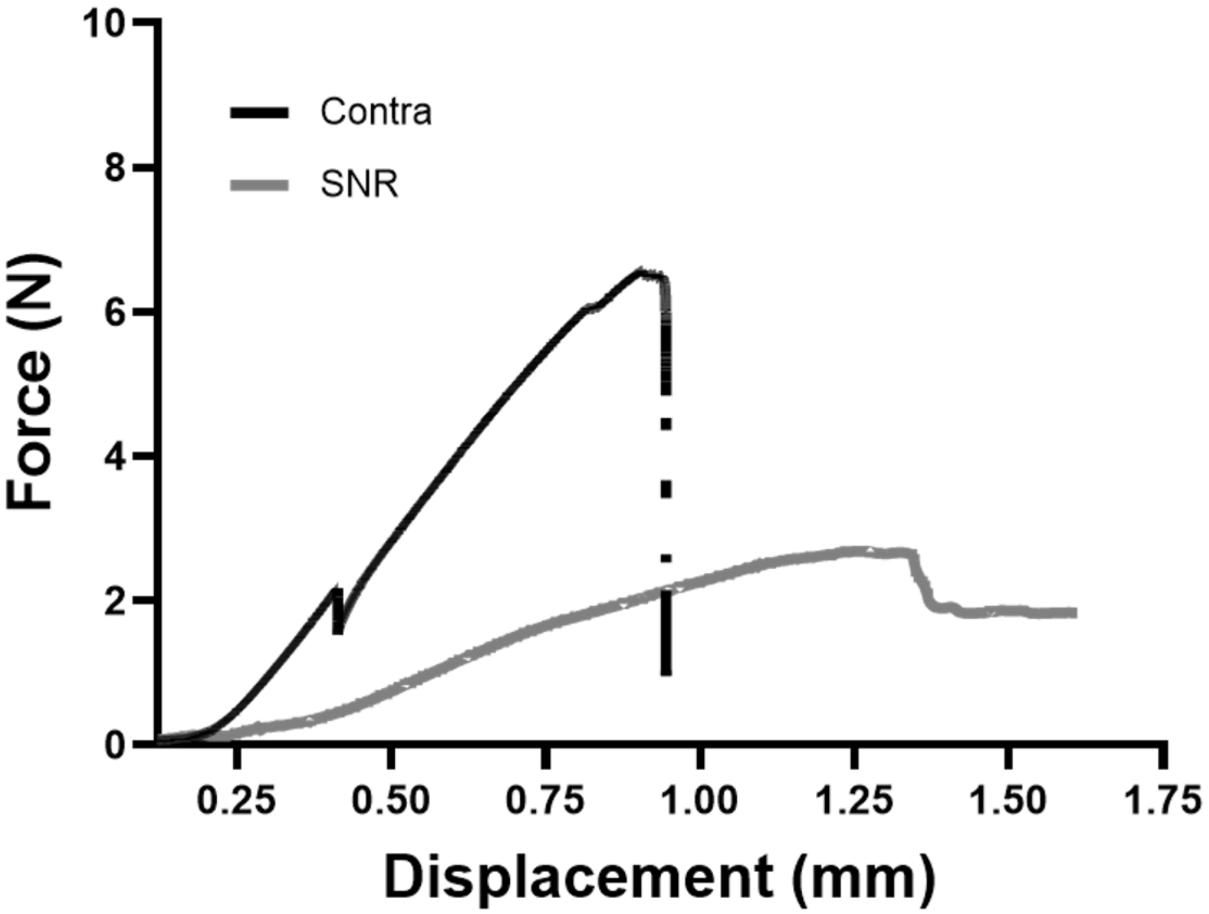

**Supplemental Figure 7: SNR alters tendon mechanical properties**. Uniaxial failure curves from representative samples from each treatment group**.**

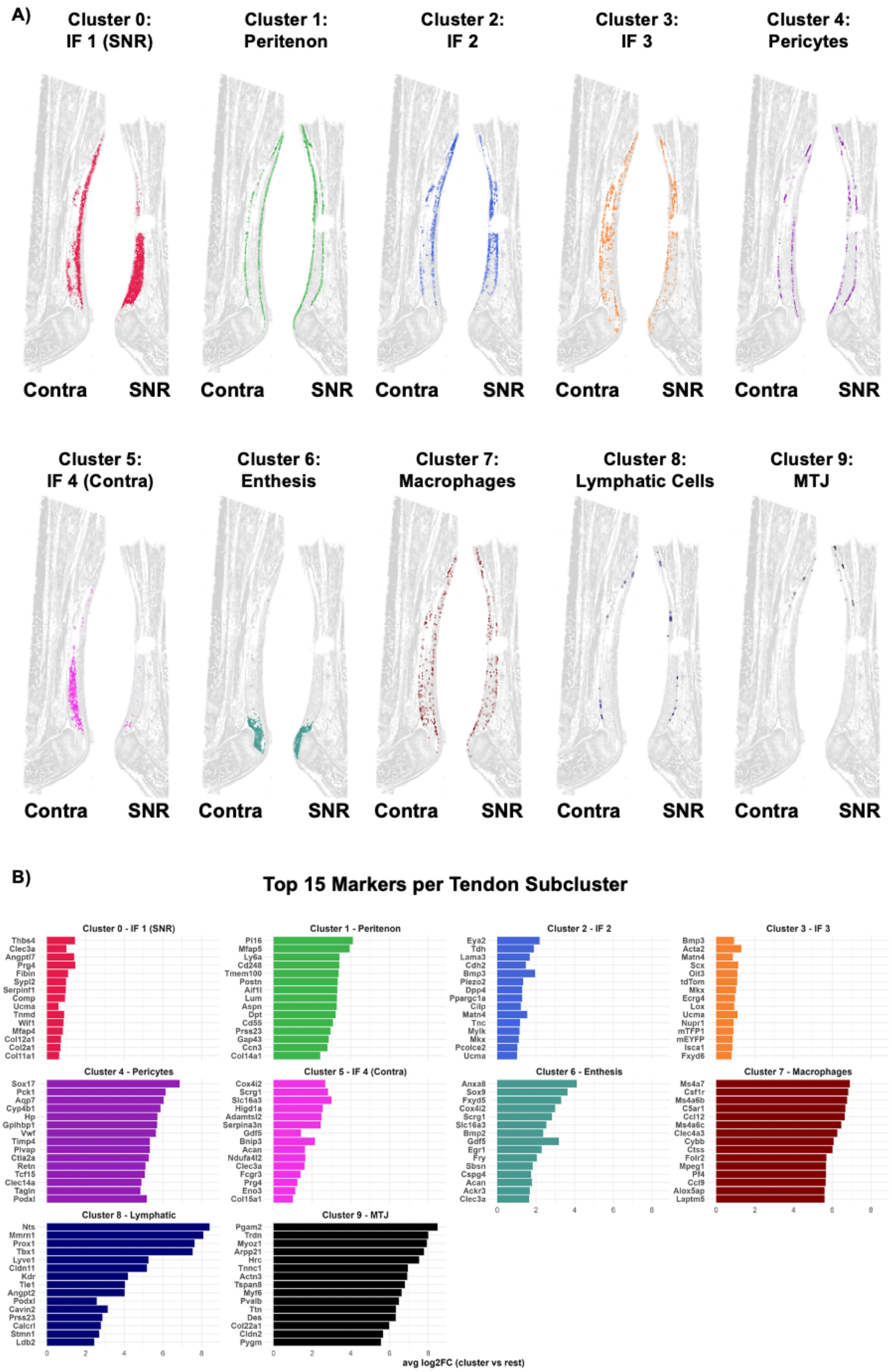
A)
Cluster 0: IF 1 (SNR)
Cluster 1: Peritenon
Cluster 2: IF 2
Cluster 3: IF 3
Cluster 4: Pericytes
Cluster 5: IF 4 (Contra)
Cluster 6: Enthesis
Cluster 7: Macrophages
Cluster 8: Lymphatic Cells
Cluster 9: MTJ
Contra
SNR
B)
Top 15 Markers per Tendon Subcluster
Cluster 0 - IF 1 (SNR)
Thbs4 Clec3a Angptl7 Prg4 Fibin Sypl2 Serpinf1 Comp Ucma Tnmd Wif1 Mfap4 Col12a1 Col2a1 Col11a1
Cluster 1 - Peritenon
Pi16 Mfap5 Ly6a Cd248 Tmem100 Postn Aif1l Lum Aspn Dpt Cd55 Prss23 Gap43 Ccn3 Col14a1
Cluster 2 - IF 2
Eya2 Tdh Lama3 Cdh2 Bmp3 Piezo2 Dpp4 Ppargc1a Cilp Matn4 Tnc Mylk Mkx Pcolce2 Ucma
Cluster 3 - IF 3
Bmp3 Acta2 Matn4 Scx Oit3 tdTom Mkx Ecrg4 Lox Ucma Nupr1 mTFP1 mEYFP Isca1 Fxyd6
Cluster 4 - Pericytes
Sox17 Pck1 Aqp7 Cyp4b1 Hp Gpihbp1 Vwf Timp4 Plvap Ctla2a Retn Tcf15 Clec14a Tagln Podxl
Cluster 5 - IF 4 (Contra)
Cox4i2 Scrg1 Slc16a3 Higd1a Adamtsl2 Serpina3n Gdf5 Bnip3 Acan Ndufa4l2 Clec3a Fcgr3 Prg4 Eno3 Col15a1
Cluster 6 - Enthesis
Anxa8 Sox9 Fxyd5 Cox4i2 Scrg1 Slc16a3 Bmp2 Gdf5 Egr1 Fry Sbsn Cspg4 Acan Ackr3 Clec3a
Cluster 7 - Macrophages
Ms4a7 Csf1r Ms4a6b C5ar1 Ccl12 Ms4a6c Clec4a3 Cybb Ctss Folr2 Mpeg1 Pf4 Ccl9 Alox5ap Laptm5
Cluster 8 - Lymphatic
Nts Mmrn1 Prox1 Tbx1 Lyve1 Cldn11 Kdr Tie1 Angpt2 Podxl Cavin2 Prss23 Calcrl Stmn1 Ldb2
Cluster 9 - MTJ
Pgam2 Trdn Myoz1 Arpp21 Hrc Tnnc1 Actn3 Tspan8 Myf6 Pvalb Ttn Des Col22a1 Cldn2 Pygm
0 2 4 6 8
avg log2FC (cluster vs rest)

C)

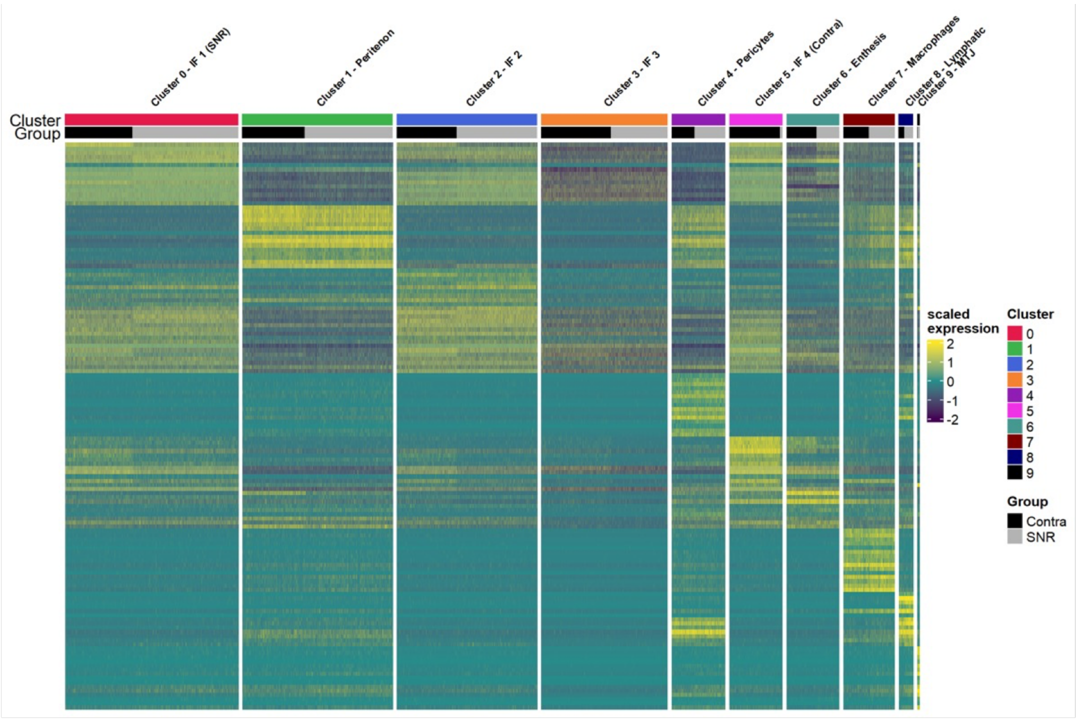

Cluster 0 - IF 1 (SNR)
Cluster 1 - Peritenon
Cluster 2 - IF 2
Cluster 3 - IF 3
Cluster 4 - Pericytes
Cluster 5 - IF 4 (Contra)
Cluster 6 - Enthesis
Cluster 7 - Macrophages
Cluster 8 - Lymphatic
Cluster 9 - MTJ
Cluster
Group
scaled expression
2
1
0
-1
-2
Cluster
0
1
2
3
4
5
6
7
8
9
Group
Contra
SNR

D)

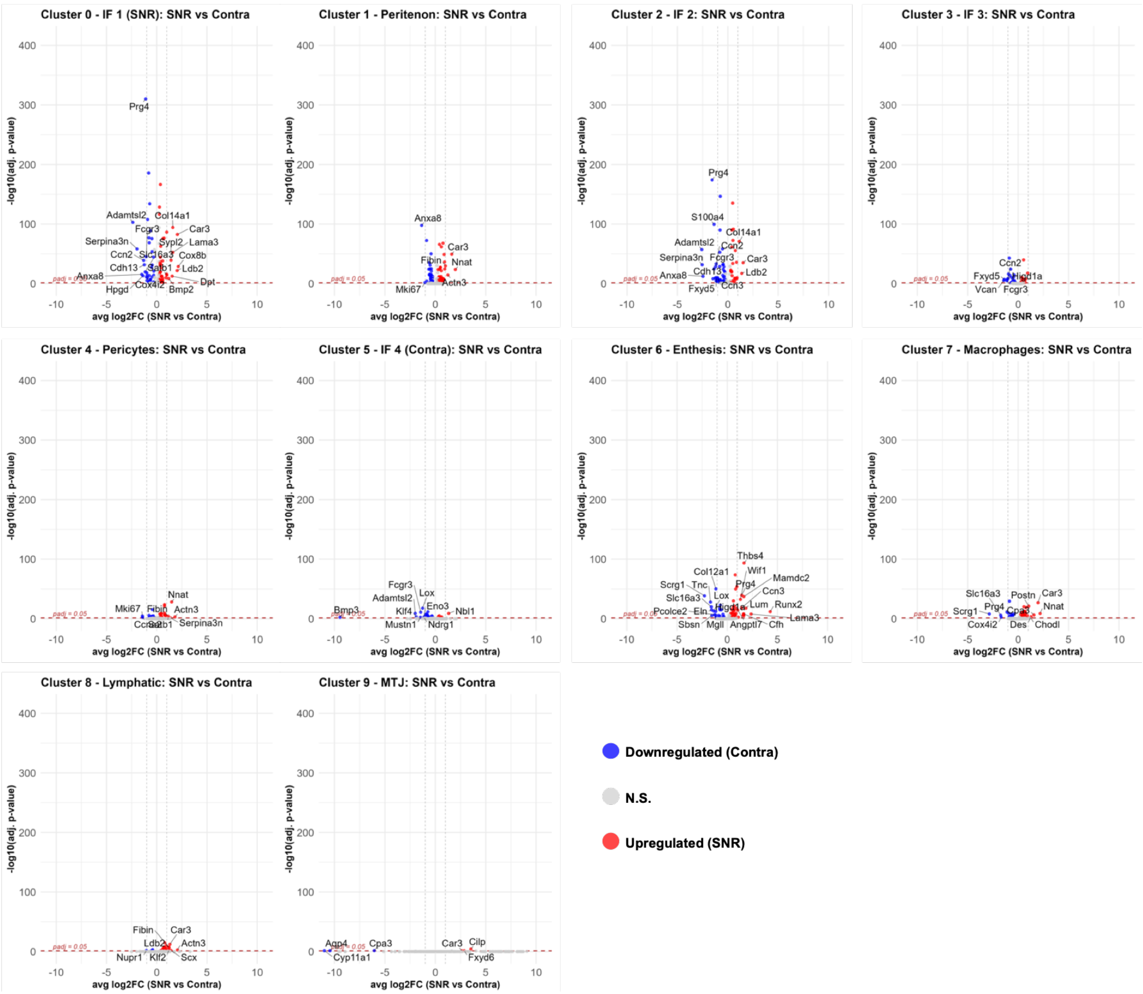

Cluster 0 - IF 1 (SNR): SNR vs Contra
Prg4
Adamtsl2
Col14a1
Fcgr3
Car3
Serpina3n
Sypl2
Lama3
Ccn2
Cox8b
Cdh13
Ldb2
Anxa8
Dpt
Hpgd
Cox4i2
Bmp2
Cluster 1 - Peritenon: SNR vs Contra
Anxa8
Car3
Fibin
Nnat
Actn3
Mki67
Cluster 2 - IF 2: SNR vs Contra
Prg4
S100a4
Col14a1
Adamtsl2
Ccn2
Serpina3n
Fcgr3
Car3
Cdh13
Ldb2
Anxa8
Fxyd5
Ccn3
Cluster 3 - IF 3: SNR vs Contra
Ccn2
Fxyd5
Hif1a
Vcan
Fcgr3
Cluster 4 - Pericytes: SNR vs Contra
Nnat
Mki67
Fibin
Actn3
Serpina3n
Cluster 5 - IF 4 (Contra): SNR vs Contra
Fcgr3
Lox
Adamtsl2
Eno3
Bmp3
Klf4
Nbl1
Mustn1
Ndrg1
Cluster 6 - Enthesis: SNR vs Contra
Thbs4
Wif1
Col12a1
Mamdc2
Scrg1
Tnc
Prg4
Ccn3
Slc16a3
Lox
Lum
Runx2
Pcolce2
Eln
Lama3
Sbsn
Mgll
Angptl7
Cfh
Cluster 7 - Macrophages: SNR vs Contra
Slc16a3
Postn
Car3
Prg4
Nnat
Scrg1
Cox4i2
Des
Chodl
Cluster 8 - Lymphatic: SNR vs Contra
Fibin
Car3
Ldb2
Actn3
Nupr1
Klf2
Scx
Cluster 9 - MTJ: SNR vs Contra
Aqp4
Cpa3
Car3
Cilp
Cyp11a1
Fxyd6
-log10(adj. p-value)
avg log2FC (SNR vs Contra)
padj = 0.05
400
300
200
100
0
-10
-5
0
5
10
Downregulated (Contra)
N.S.
Upregulated (SNR)

E)

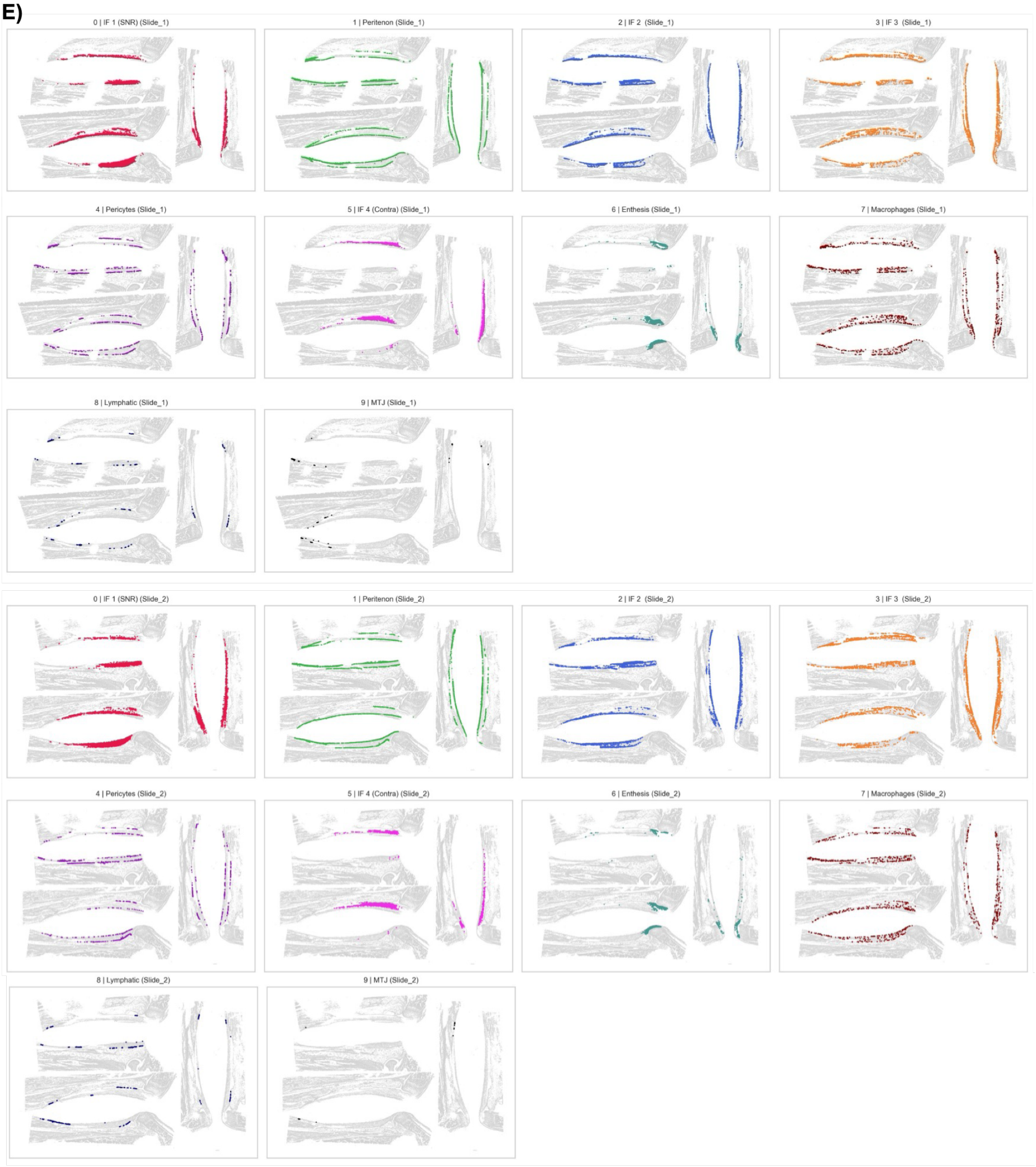
0 | IF 1 (SNR) (Slide_1)
1 | Peritenon (Slide_1)
2 | IF 2 (Slide_1)
3 | IF 3 (Slide_1)
4 | Pericytes (Slide_1)
5 | IF 4 (Contra) (Slide_1)
6 | Enthesis (Slide_1)
7 | Macrophages (Slide_1)
8 | Lymphatic (Slide_1)
9 | MTJ (Slide_1)
0 | IF 1 (SNR) (Slide_2)
1 | Peritenon (Slide_2)
2 | IF 2 (Slide_2)
3 | IF 3 (Slide_2)
4 | Pericytes (Slide_2)
5 | IF 4 (Contra) (Slide_2)
6 | Enthesis (Slide_2)
7 | Macrophages (Slide_2)
8 | Lymphatic (Slide_2)
9 | MTJ (Slide_2)

F)

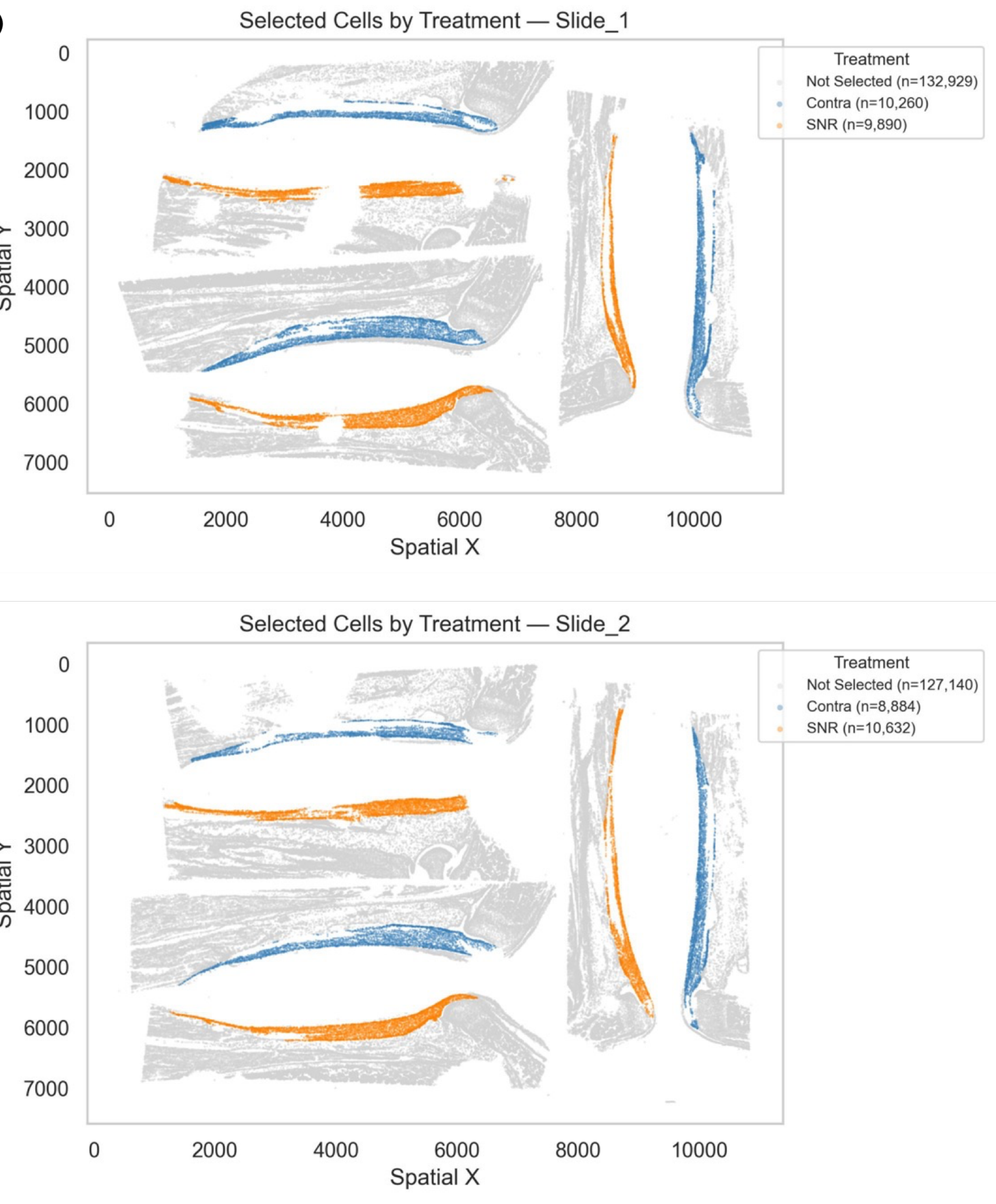

**Supplemental Figure 8. Xenium analysis of Achilles tendon cell cluster identity, spatial distribution, and cluster abundance.** A) Spatial mapping of the 10 Leiden clusters illustrating the anatomical distribution of each transcriptionally defined cell population within the tendon sections. B) Top-ranked differentially expressed genes for each cluster based on $\log_2$ fold change. C) Heatmap of scaled gene expression across tendon subclusters and treatments, demonstrating cluster-specific expression patterns. D) Volcano plots showing differential gene expression between SNR and contralateral cells within each subclusters. Positive $\log_2$ fold-change values indicate genes enriched in SNR (red), whereas negative values indicate genes enriched in contralateral cells (blue). E) Spatial distribution of the 10 tendon subclusters across all tissue sections from slide 1 and slide 2. F) Spatial mapping of selected tendon cells by treatment across Slides 1 and 2, showing the anatomical localization of contralateral (blue) and SNR (orange) cells within the tendon regions used for subcluster analysis (n = 3).

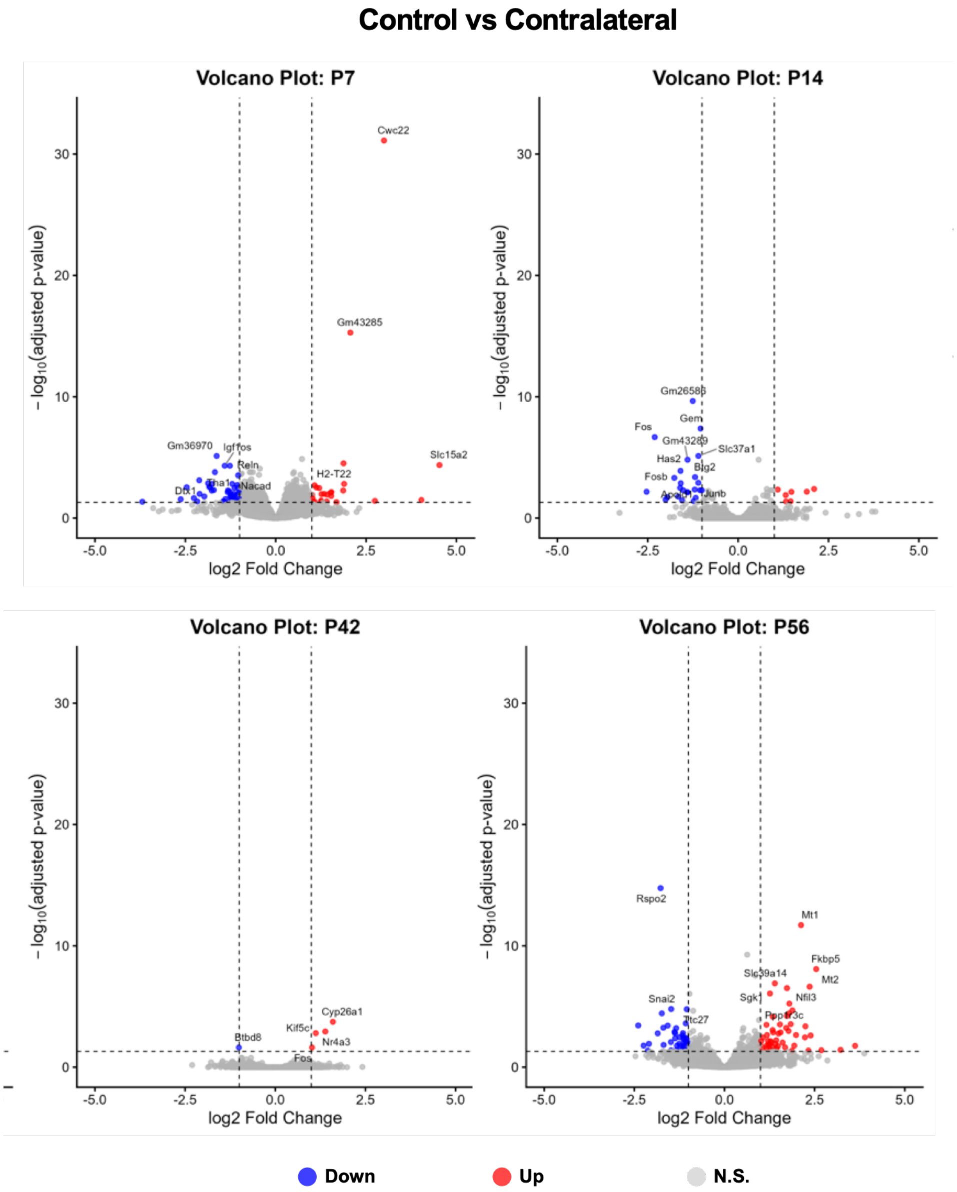

**Supplemental Figure 9: Contralateral limbs are comparable to uninjured controls across transcriptional analyses.** Bulk RNA-sequencing analysis across developmental timepoints (P7, P14, P42, P56) revealed minimal differential gene expression between control and contralateral tendons, as shown by volcano plots with few significantly up- or downregulated genes (n = 4-5).

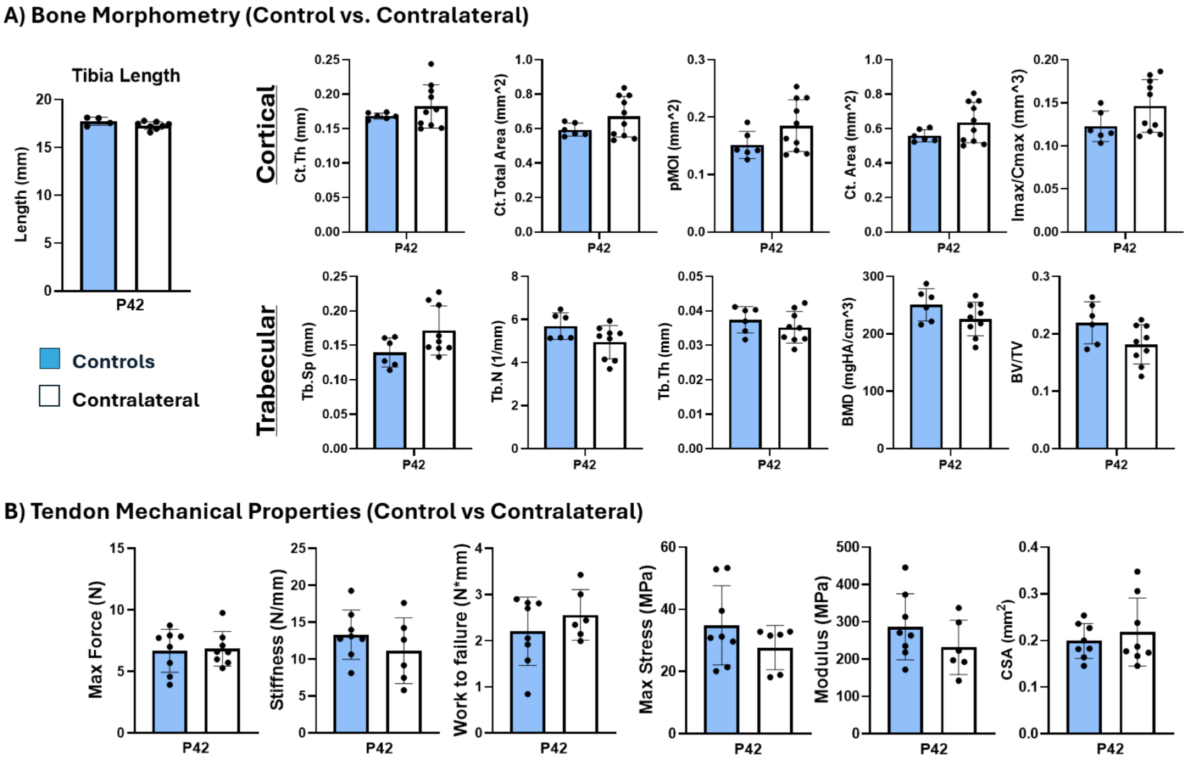

**Supplemental Figure 10: Contralateral limbs are comparable to uninjured controls across bone parameters and tendon mechanical properties.** A) Bone analysis demonstrated no significant difference between contralateral bone parameters compared to non-injured controls following SNR ($n = 4$-$8$, $p < 0.05$). B) Mechanical testing of Achilles tendons showed comparable functional properties between groups, including maximum force, stiffness, stress, and modulus ($n = 6$-$8$, $p < 0.05$).

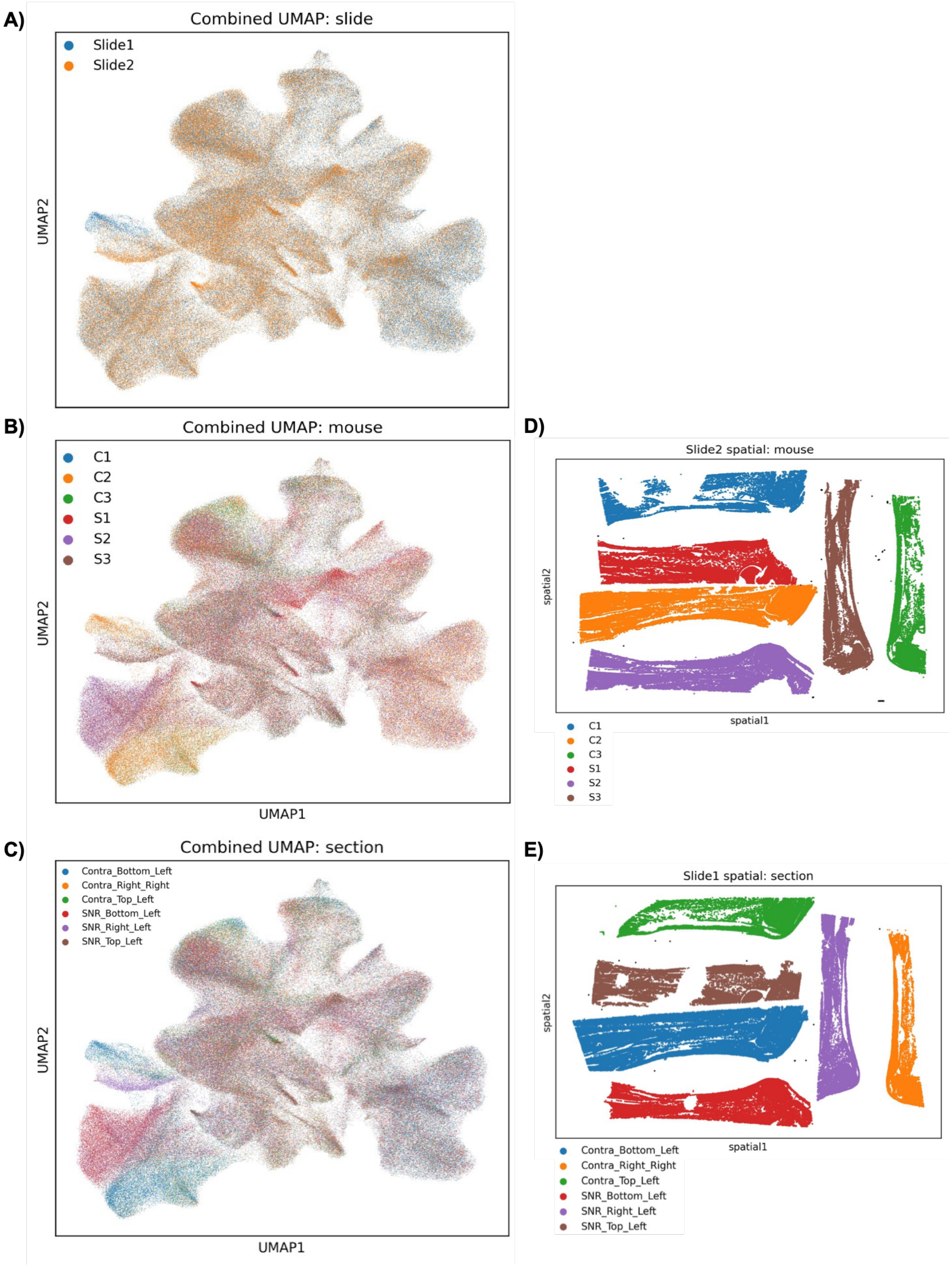

**Supplemental Figure 11: Xenium dataset across slides, animals, and tissue sections.** A) Combined UMAP colored by Xenium slide, demonstrating overlap of cells from Slides 1 and 2 following integration. B) Combined UMAP colored by individual mouse and treatment, where C1–C3 represent the contralateral limbs and S1–S3 represent the SNR limbs from mice 1–3, respectively. C) Combined UMAP colored by individual tissue section. D) Spatial distribution of cells from Slide 2 colored by mouse and treatment. E) Spatial distribution of cells from Slide 1 colored by tissue section.

**Supplemental Table 2:** 10x Xenium Spatial Transcriptomics probes for 379 genes in the "Mouse Tissue Atlassing" panel, in addition to 100 custom genes.

| Tissues Atlas Panel | | | | | | Custom Genes | |
|---|---|---|---|---|---|---|---|
| Sox17 | Riiad1 | Rho | Rbp2 | Myoz1 | Cp | Acta2 | Prg4 |
| Il1r2 | Celf3 | Fxyd4 | Arpp21 | Mustn1 | Cpa3 | Adamts17 | Scrg1 |
| Pou3f3 | Selenbp1 | Mfap5 | Iyd | Tnnc1 | Tmem212 | Adamtsl2 | Scx |
| Ecrg4 | Ctss | Clec4a3 | Trdn | Anxa8 | Cldn11 | Angptl7 | Slc16a3 |
| Cavin2 | Fmo5 | Vwf | Pln | Fam25c | Sox2 | Bmp3 | Slc1a3 |
| Ctla4 | Hmgcs2 | Kcna1 | Oit3 | 2610528A11Rik | Tm4sf4 | Bnip3 | Slc8a1 |
| Cxcr2 | Hsd3b4 | Klra8 | Gamt | Sftpd | Gria2 | Ccn1 | Spon2 |
| Des | Sypl2 | Plbd1 | Timp3 | Ctsg | Cd5l | Ccn2 | Sun1 |
| Scg2 | 5330417C22Rik | Ldhb | Ascl1 | Slc25a37 | Muc1 | Ccna2 | Sun2 |
| Tm4sf20 | F3 | C5ar1 | Lum | Epsti1 | S100a14 | Cd248 | Tdh |
| Ackr3 | Myoz2 | Pnmal2 | Mgat4c | Ccn3 | S100a4 | Cd44 | Tgfb1 |
| Serpinb5 | Uox | Pglyrp1 | Nts | Ndrg1 | Lor | Cd55 | Tgfb2 |
| Serpinb1 | Calb1 | Apoc2 | Myf6 | Ly6h | Ivl | Cdca8 | Tgfbr2 |
| Tfcp2l1 | Atp6v0d2 | Apoc4 | Tspan8 | Gpihbp1 | Crct1 | Cdh11 | Tgfbr3 |
| Marco | Aqp7 | Cnfn | Wif1 | Pvalb | Lce1m | Cdh13 | Thbs1 |
| Ctse | Aqp3 | Prx | Ndufa4l2 | Ncf4 | Them5 | Cdh2 | Tnc |
| Cfh | 1110017D15Rik | Aplp1 | Pgam2 | Rac2 | Bgn | Cdk1 | Tnmd |
| Dpt | Aldh1b1 | Upk1a | Abca13 | Upk3a | Plp1 | Cdt1 | Top2a |
| Gpa33 | Hemgn | Tmem147 | Gabrp | Mlc1 | Asz1 | Cilp | Tppp3 |
| Rgs5 | Ambp | Sbsn | Ltc4s | Col2a1 | Podxl | Clec3a | Vcan |
| Cenpf | Rab3b | Fxyd1 | Fam183b | Aqp5 | Ptn | Col11a1 | Wwtr1 |
| Prox1 | Cyp4b1 | Fxyd3 | Rasd1 | Mettl7a2 | Tmem213 | Col12a1 | Yap1 |
| Ucma | Tie1 | Slc17a7 | Mfap4 | Krt79 | Try10 | Col14a1 | mEYFP |
| Itga8 | Guca2b | Hrc | Pmp22 | Krt8 | Prss3 | Col15a1 | mTFP1 |
| Cubn | Csf3r | 0610005C13Rik | Cldn7 | Igfbp6 | Aqp1 | Col22a1 | tdTom |
| Pax8 | Gjb4 | Mrgpra2a | 6330403K07Rik | Ppp1r1a | Nap1l5 | Col5a1 | Mmp13 |
| Fcnb | Laptm5 | A330076H08Ri | Serpinf1 | AU021092 | Mmrn1 | Cox4i2 | Mmp14 |
| Aif1l | Stmn1 | Isg20 | Ccl12 | Myh11 | Cd8a | Csf1 | Mmp2 |
| Eng | Rhd | Acan | Ccl9 | Tbx1 | Reg3g | Csf1r | Myh10 |
| Hc | Nbl1 | Rhcg | Mpo | Lamp3 | Clec4f | Cspg4 | Myh9 |
| Lhx2 | Clcnka | Nox4 | Tmem100 | Hrg | Atp6v1b1 | Dcn | Nfatc2 |
| Tnfaip6 | Clcnkb | Prss23 | B4galnt2 | Kng2 | Asprv1 | Dlc1 | Ooep |
| Abcb11 | Rbp7 | Omp | Krt13 | Mylk | Mgll | Dpp4 | Pdgfa |
| Lrp2 | Cd36 | Folr2 | Krt19 | Stfa2l1 | Chl1 | Egr1 | Piezo1 |
| Gad1 | Slc5a1 | Hpx | Krt16 | Cstdc4 | Timp4 | Eln | Piezo2 |
| Ttn | Htra3 | Lyve1 | Hap1 | Upk1b | Cldn2 | Eno3 | Plcb1 |
| Neurod1 | Bst1 | Gp2 | Slc4a1 | Gap43 | Tubb3 | Epha4 | Postn |
| Calcrl | Ppargc1a | Nupr1 | Higd1b | Ccdc80 | Mmp8 | Esr1 | |
| Prg2 | Sod3 | Prap1 | Gfap | Retnla | Cnn1 | Eya2 | |
| Gm13889 | Slc34a2 | Cox8b | Wnt3 | Col8a1 | Kcnj1 | Fap | |
| Fibin | Kdr | Ifitm6 | Kcnj16 | Chodl | Trim29 | Fcgr3 | |
| Scg5 | Slc4a4 | Cdhr5 | Sox9 | Dcpp2 | Ccdc153 | Fn1 | |
| Gatm | Gc | Sct | Cd300lf | Pi16 | Hmbs | Fry | |
| Slc27a2 | Ppbp | Retn | Sdc1 | Tff2 | Ttc36 | Fxyd5 | |
| Prom2 | Pf4 | Angpt2 | Vsnl1 | Psmb8 | Cd3d | Gdf5 | |
| Chgb | Areg | Sfrp1 | Rsad2 | Ager | Fxyd6 | Gja1 | |
| Bmp2 | Mapk10 | Hpgd | Colec11 | Ly6g6c | Tagln | Gmnn | |
| Snap25 | Sez6l | Aadat | Myt1l | Lst1 | Cryab | Higd1a | |
| Otor | Dao | Comp | Sostdc1 | Sfta2 | Cyp11a1 | Ifrd1 | |
| Insm1 | Cux2 | Tmem59l | Etv1 | Runx2 | Anxa2 | Igfbp3 | |
| Tcf15 | Cldn4 | Plvap | Scin | Satb1 | Scg3 | Itgb5 | |
| Myl9 | Cldn3 | Mylk3 | Coch | C3 | Pcolce2 | Itih5 | |
| Nnat | Upk3b | Capns2 | Clec14a | Epcam | Rbp1 | Kcnq5 | |
| Ube2c | Actl6b | Cdh16 | Dbpht2 | Aqp4 | Ace2 | Klf2 | |
| Mmp9 | Alox5ap | Hsd11b2 | Gpx2 | Dsc3 | Serpina10 | Klf4 | |
| Pck1 | Kl | Hp | Slc10a1 | Scgb3a2 | Serpina3n | Lama3 | |
| Car3 | Gng11 | Tat | Vsx2 | Gm94 | Slc25a47 | Ldb2 | |
| Car2 | Cav1 | Dynlrb2 | Fbln5 | Cdx1 | Ptprn2 | Lmna | |
| Calm4 | Isca1 | Pygm | Mpeg1 | Fech | Cndp2 | Lox | |
| Calm5 | Ctla2a | Slc22a8 | Mamdc2 | Lipg | Ndufs8 | Ly6a | |
| Gpx6 | Thbs4 | Slc22a6 | Col17a1 | Slc14a1 | Actn3 | Matn4 | |

| | | | | | | |
|---|---|---|---|---|---|---|
| Dcdc2a | Emb | Ms4a7 | Cybb | Slc14a2 | Cplx2 | Mki67 |
| F13a1 | Dnase1l3 | Ms4a6c | Aspn | Oit1 | Ms4a6b | Mkx |
| | | | | | Asb9 | |
